# Oligomerization-dependent and synergistic regulation of Cdc42 GTPase cycling by a GEF and a GAP

**DOI:** 10.1101/2023.06.26.546500

**Authors:** Sophie Tschirpke, Werner K-G. Daalman, Frank van Opstal, Liedewij Laan

## Abstract

Cell polarity is a crucial biological process essential for cell division, directed growth, and motility. In *Saccharomyces cerevisiae*, polarity establishment centers around the small Rho-type GTPase Cdc42, which cycles between GTP-bound and GDP-bound states, regulated by GEFs like Cdc24 and GAPs such as Rga2. To dissect the dynamic regulation of Cdc42, we employed *in vitro* GTPase assays, revealing inverse concentration-dependent profiles for Cdc24 and Rga2: with increasing concentration, Cdc24’s GEF activity is non-linear and oligomerization-dependent, which is possibly linked to the relief of its self-inhibition. In contrast, Rga2’s GAP activity saturates, likely due to self-inhibition upon oligomerization. Together, Cdc24 and Rga2 exhibit a strong synergy driven by weak Cdc24 - Rga2 binding. We propose that the synergy stems from Cdc24 alleviating the self-inhibition of oligomeric Rga2. We believe this synergy contributes to efficient regulation of Cdc42’s GTPase cycle over a wide range of cycling rates, enabling cells to resourcefully establish polarity. As Cdc42 is highly conserved among eukaryotes, we propose the GEF-GAP synergy to be a general regulatory property in other eukaryotes.

## Introduction

Cells require robust, yet adaptable, functioning to survive in an ever-changing environment. One of such functionalities is cell polarity, which is essential for processes such as cell division, directed growth and secretion, and motility [Vendel et al., 2019]. A well-studied system for polarity establishment is that of *Saccharomyces cerevisiae*: here the cell division control protein Cdc42 exits the cytoplasm and accumulates in one spot on the cell membrane, marking the site of bud emergence (Fig. 1a) [Nelson, 2003, Etienne-Manneville, 2004, Thompson, 2013, Diepeveen et al., 2018, Chiou et al., 2017, Martin, 2015]. Cdc42 is a highly conserved small Rho-type GTPase, who’s activity involves cycling between a GTP-bound state and a GDP-bound state, a process tightly controlled by guanine nucleotide exchange factors (GEFs), such as Cdc24 and Bud3, and GTPase activating proteins (GAPs), such as Rga1, Rga2, Bem2, and Bem3 [Park and Bi, 2007, Martin, 2015] (Fig. 1b).

**Figure 1.**
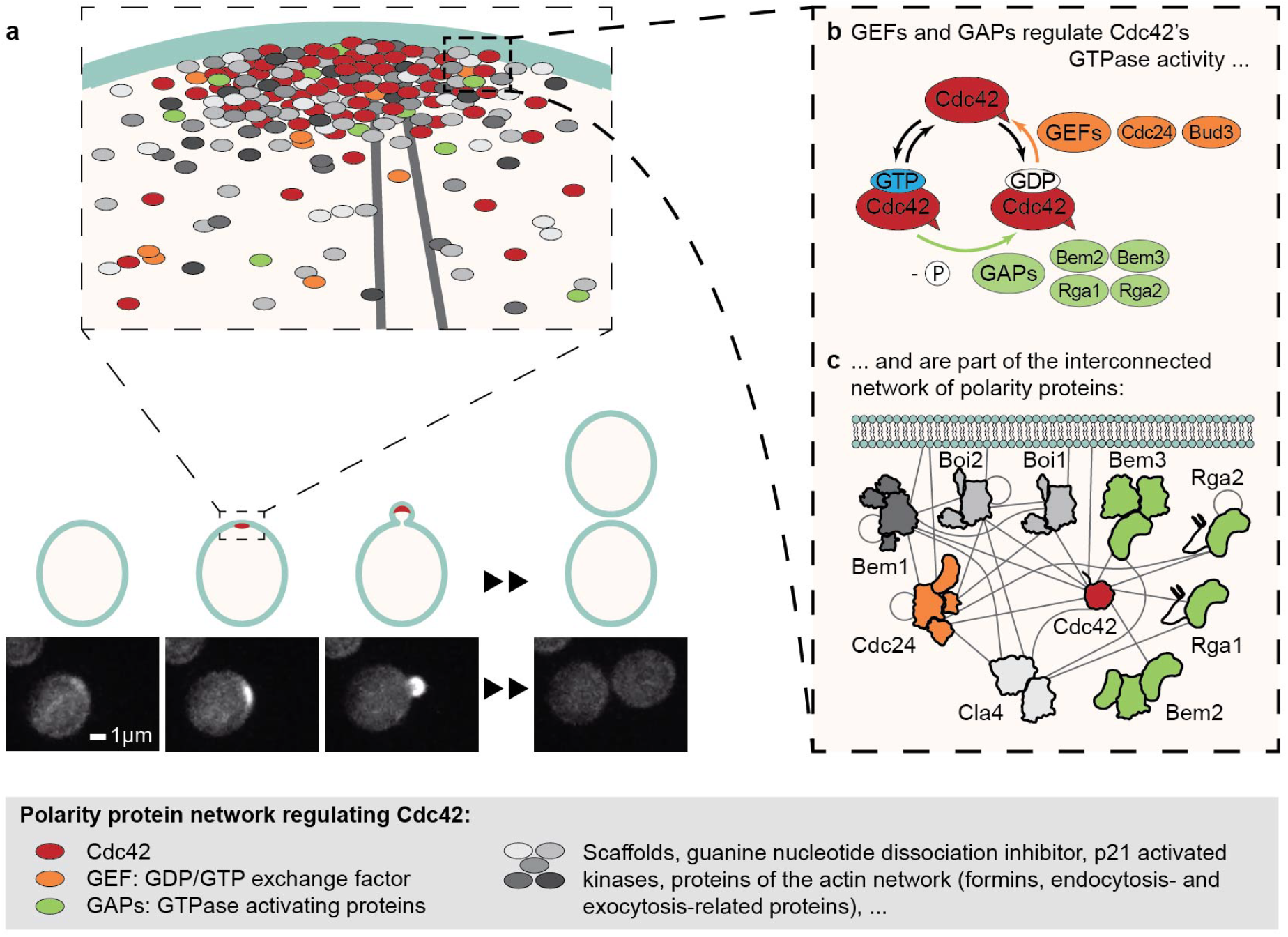
A complex network of polarity proteins regulates Cdc42 activity *in vivo* to establish cell polarity. (a) The accumulation of the small Rho-GTPase Cdc42 in one spot on the cell membrane establishes cellular polarity, initiating cell division of *Saccharomyces cerevisiae*. Polarity establishment is driven by interactions between Cdc42 and polarity proteins of an intricate network. Cdc42 is shown in red in the cartoon and in white in life cell microscopy images (bottom). (b) The polarity protein groups of GEFs and GAPs regulate Cdc42 GTPase cycling, a process required for its functioning in polarity establishment. (c) GEFs and GAPs are part of the interconnected and complex polarity protein network. This cartoon shows a subset of polarity proteins and grey lines depict experimentally determined (direct or indirect) interactions. Protein subdomains roughly correspond to domain size but do not accurately depict the domain structure.

A thorough understanding of Cdc42 GTPase activity and its regulation is crucial, as Cdc42 GTPase cycling is required for polarity establishment and cells in which it has been impaired fail to polarize [Wedlich-Soldner et al., 2004]. Cdc42 GTPase activity and its role in polarity establishment has been studied both *in vivo* [Smith et al., 2013, Lee et al., 2015, Sartorel et al., 2018] and *in vitro*. However, interpreting *in vivo* studies can be challenging due to the complexity within the regulatory network surrounding Cdc42. For instance, several proteins belong to the same group of regulators, leading to redundancy within the network. Further, many polarity proteins also interact with a variety of other polarity proteins, resulting in a dynamic assembly of loosely interacting proteins at the polarity spot [Gao et al., 2011, Daalman et al., 2020] (Fig. 1c). *In vitro* studies, where interactions can be probed in a highly controlled environment, are still comparatively scarce, focus on a small aspect of the system, and did not include interactions between different full-length regulatory proteins [Zheng et al., 1993, Zheng et al., 1994, Zheng et al., 1995, Zhang et al., 1997, Zhang and Zheng, 1998, Zhang et al., 1999, Zhang et al., 2001, Smith et al., 2002, Das et al., 2012, Johnson et al., 2009, Johnson et al., 2012, Golding et al., 2019]. Therefore they neglect effects present in multi-protein systems as occur in the dynamic and interconnected structure of the polarity network *in vivo* (Fig. 1c). A recent study confirms this notion, revealing that the scaffold Bem1 enhances the GEF activity of Cdc24 [Rapali et al., 2017].

To enhance our knowledge of Cdc42 regulation in *S. cerevisiae*, we employ a bulk *in vitro* GTPase assay to study properties of the entire Cdc42 GTPase cycle and its regulation by the GEF Cdc24 and GAP Rga2. These proteins are interesting study targets; Cdc24 is an essential protein and one of the two GEFs present at the polarity site [Daalman et al., 2020], and Rga2 has not yet been characterized as full-length protein [Smith et al., 2002]. Both proteins have the potential to oligomerize [Mionnet et al., 2008, Tarassov et al., 2008, Schlecht et al., 2012], a property which may enhance polarity establishment [Lang and Munro, 2022] and could tune their activity in a dosage-dependent fashion. Lastly, Cdc24 and Rga2 regulate distinct steps of the Cdc42 GTPase cycle and interact *in vivo* (although it is unclear whether their interaction is direct or indirect) [McCusker et al., 2007, Breitkreutz et al., 2010, Chollet et al., 2020]. In combination Cdc24 and Rga2 could synergize, inhibit each other, or have no interplay. Such a GEF - GAP interaction is theorized to play a significant role in G-protein signaling [Ross, 2008].

Our data reveals distinct concentration-and oligomerization-dependent profiles for the GEF and GAP: Specifically, Cdc24 stimulates Cdc42 GTPase activity in a non-linear fashion, a process that is linked to its oligomerization and may relieve its self-inhibition. Rga2 also forms oligomeric structures, driven by its C-terminal GAP domain. Interestingly, the GAP activity of full-length Rga2, unlike that of its isolated GAP domain, saturates at an approximate 1:1 ratio of Rga2 to Cdc42. This saturation may be linked to self-inhibition upon oligomerization. In combination, Cdc24 and Rga2 exhibit a protein-specific GEF-GAP synergy, which appears to arise from weak Cdc24-Rga2 binding. We speculate that it arises from the relieve of oligomeric Rga2’s self-inhibition through Cdc24.

## Results

### Cdc42 GTPase activity can be reconstituted *in vitro*

We used the GTPase-Glo™ assay (Promega) to examine the entire GTPase cycle of Cdc42. This allowed us to study the interplay of regulators acting on different steps of the GTPase cycle. In each assay, Cdc42, alone or in combination with one or multiple effector proteins, is mixed with GTP and incubated at 30^°^C for 1-1.5 h for GTPase cycling to occur. Then the reaction is stopped through transforming the remaining GTP to luminescent ATP. The luminescent signal of each reaction mixture is measured (Fig. 2). The decrease of the GTP concentration during GTPase cycling is well fitted by an exponential model (S1 Fig. 1):

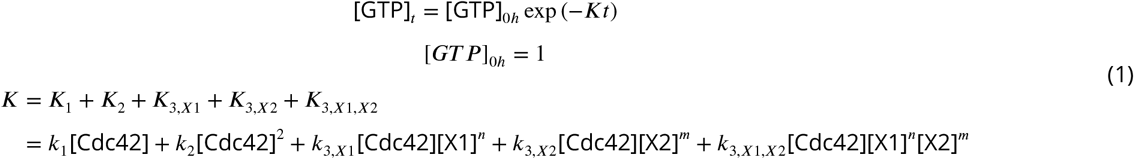

where *K* refers for the overall GTP hydrolysis rate, *X*1 and *X*2 are effector proteins, such as GEFs and GAPs, and *n* and *m* are natural numbers. From these fits the GTPase cycling rates *k* are determined [Tschirpke et al., 2024].

**Figure 2.**
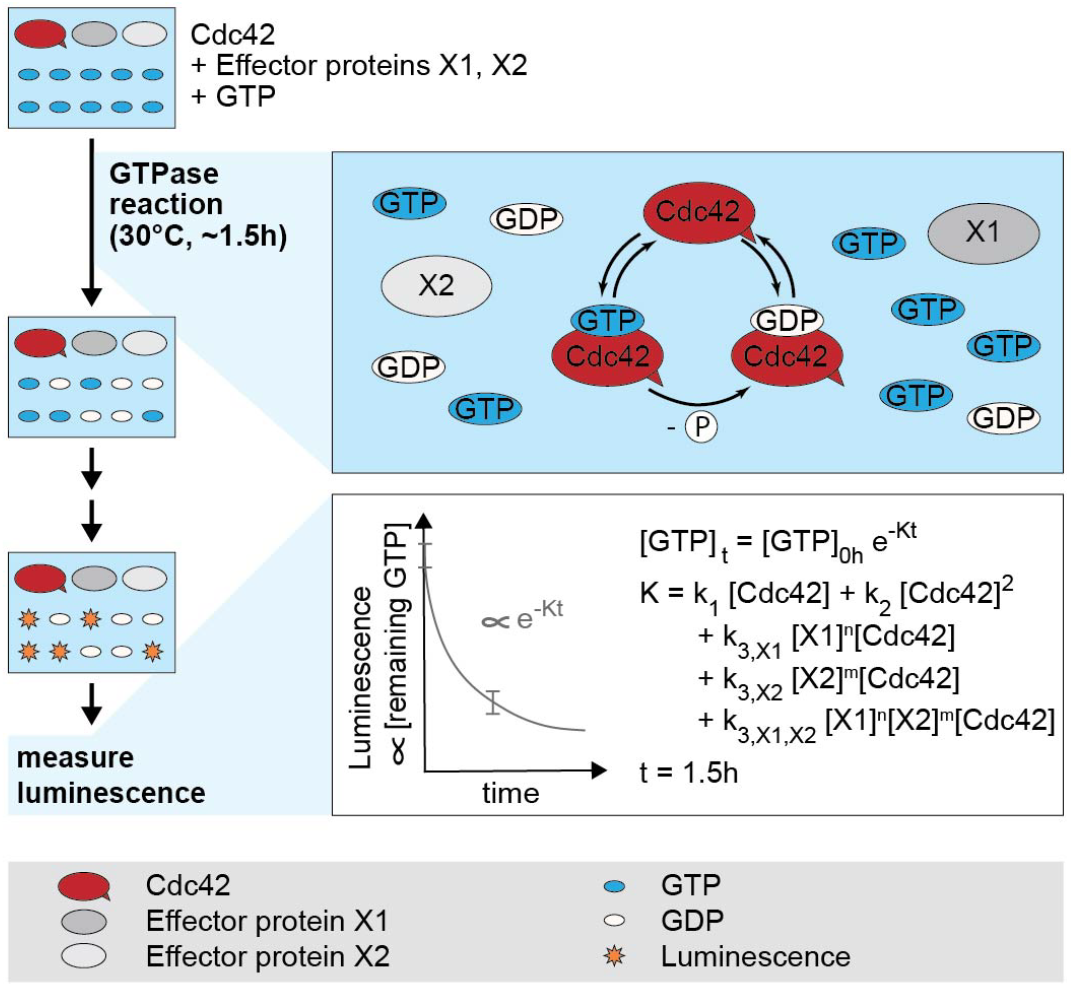
Reconstitution of Cdc42 GTPase activity regulation *in vitro*. Bulk *in vitro* GTPase assays (GTPase-Glo™ assay, Promega) allow to study the GTPase cycling of Cdc42. Cdc42, alone or in combination with effector proteins, is incubated with GTP at 30^°^C for *∼*1.5 h in which GTPase cycling occurs. After two processing steps, the amount of remaining GTP, measured as luminescence, is assessed. GTP hydrolysis cycling rates can be extracted by fitting the data with an exponential model (S1) [Tschirpke et al., 2024].

We first analyzed the GTPase activity of Cdc42 alone (Fig. 3a). To describe Cdc42 GTPase cycling, we included terms that depend linearly (*K*_1_) and quadratically (*K*_2_) on the Cdc42 concentration, the latter representing any possible effects due to cooperativity from dimeric Cdc42. While *in vivo* data suggests that *S. cerevisiae* Cdc42 does not dimerize [Kang et al., 2010], our *in vitro* GTPase assays revealed variability in Cdc42 behavior, with some Cdc42 constructs exhibiting quadratic rate increases and others showed linear ones [Tschirpke et al., 2023b]. As our goal was to explore Cdc42 - effector interactions, we used the general phenomenological description shown in Eq. 1 to fit the Cdc42 data (Fig. 3, Tab. 1). We remain cautious from deriving conclusions from this data on Cdc42 dimerization.

**Table 1.**
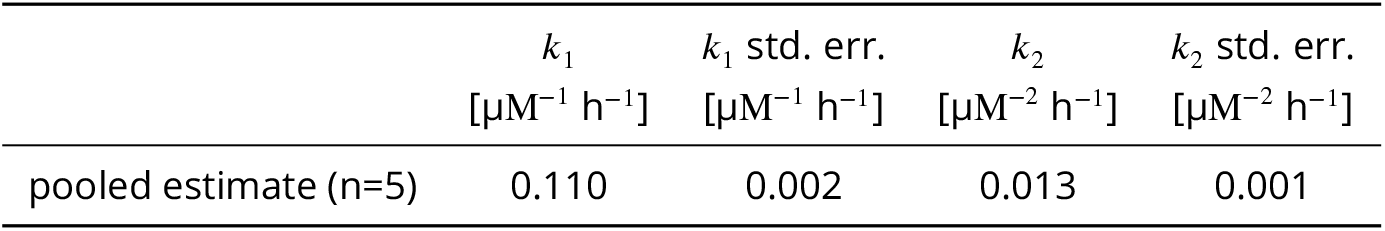
GTP hydrolysis cycling rates *k*_1_ and *k*_2_ of Cdc42.

| | $k_1$ | $k_1$ std. err. | $k_2$ | $k_2$ std. err. |
| --- | --- | --- | --- | --- |
| | $[\mu\text{M}^{-1} \text{h}^{-1}]$ | $[\mu\text{M}^{-1} \text{h}^{-1}]$ | $[\mu\text{M}^{-2} \text{h}^{-1}]$ | $[\mu\text{M}^{-2} \text{h}^{-1}]$ |
| pooled estimate (n=5) | 0.110 | 0.002 | 0.013 | 0.001 |

**Figure 3.**
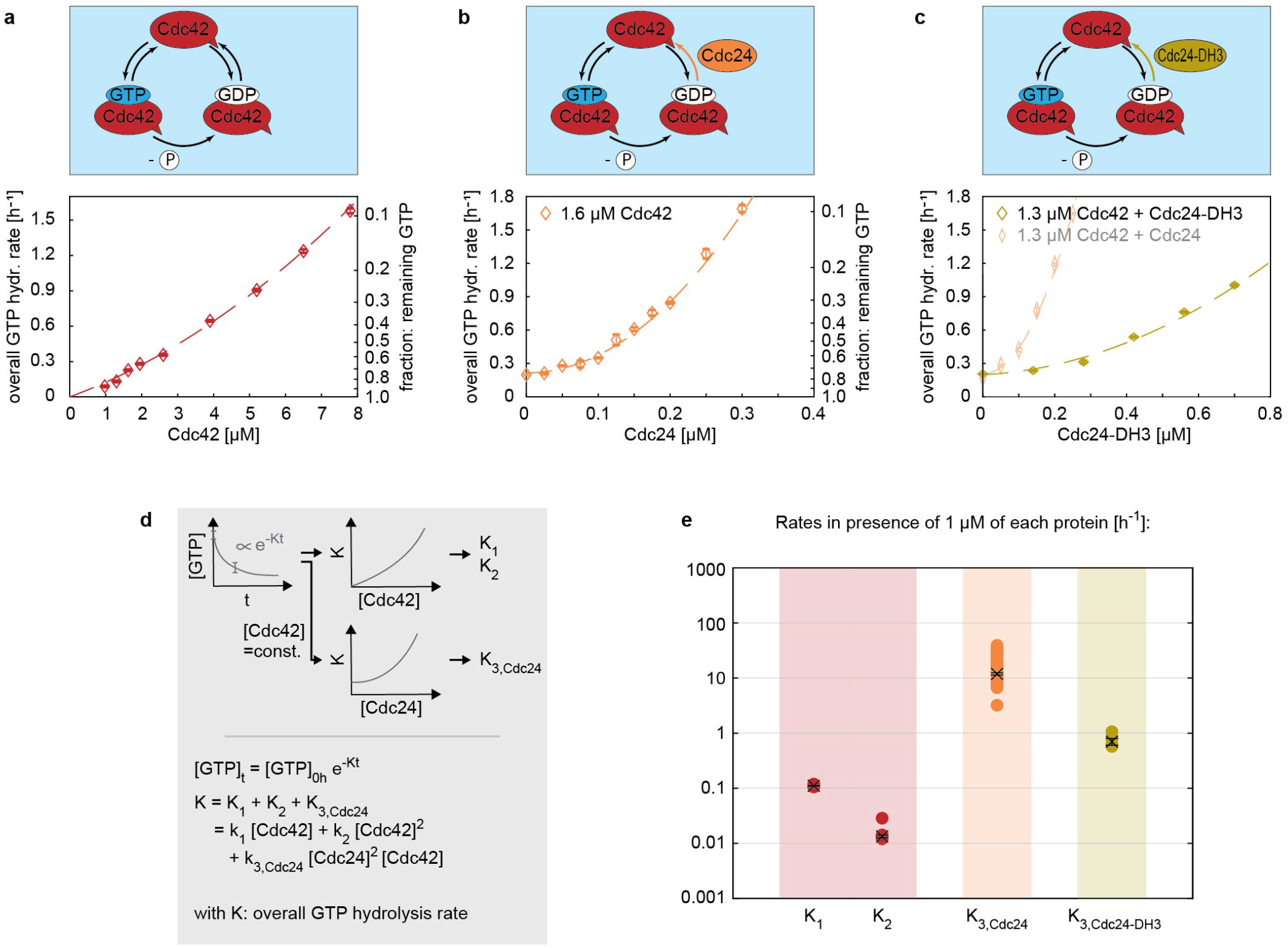
The strong GEF activity of Cdc24 requires oligomerization. (a) The overall GTP hydrolysis rate *K* of the GTPase Cdc42 in absence of any effector proteins. (b) Cdc24 boosts the GTPase activity of Cdc42 in a non-linear fashion and is active even at sub-µM concentrations. (c) Cdc24-DH3, a Cdc24 mutant with reduced oligomerization capacity, exhibits a weaker GEF activity. (d) Illustration of the data processing and fitting model. (e) Summary of the rates *K*. The values shown refer to the rate values in presence of 1 µM of each protein, e.g. ‘*K*_3,*Cdc*24_’ refers to ‘*k*_3,*Cdc*24_[Cdc24]^2^[Cdc42]’ with [Cdc42]=[Cdc24]=1 µM. Crosses with error bars represent the weighted mean and the standard error of the mean (S1), filled circles show individual measurements.

### The GEF activity of Cdc24 increases upon oligomerization

We next investigated how Cdc24 affects Cdc42 GTPase activity. Cdc24 is a known GEF, increasing the speed of the GDP release step. In agreement with previous studies [Rapali et al., 2017], sub-µM concentrations of Cdc24 substantially boost Cdc42’s GTPase activity; the rate-contribution of Cdc24 is two orders of magnitude greater than those of Cdc42 alone (Fig. 3e, Tab. 2). We find that Cdc24’s effect increases non-linearly with its concentration (Fig. 3b), which could be explained by Cdc24 di-or oligomerization: Cdc24 has the capability to oligomerize via its DH domain [Mionnet et al., 2008]^1^. Dimers and oligomers could have an increased GEF activity through releasing Cdc24 from its self-inhibited state [Shimada et al., 2004]. With increasing Cdc24 concentration the amount of Cdc24 oligomers increases, resulting in a non-linear increase of the overall GTP hydrolysis rate *K*.

**Table 2.** Interaction rates *k*_3,*X*_ of Cdc42 - effector protein mixtures. *: unit in case of X=Cdc24: [µM^−3^ h^−1^].

| | Effector protein X | $k_{3,X}$<br>[$\mu M^{-2} h^{-1}$] * | $k_{3,X}$<br>std. err. |
| --- | --- | --- | --- |
| pooled est. (n=34) | Cdc24 | 11.887 | 0.594 |
| pooled est. (n=4) | Cdc24-DH3 | 0.702 | 0.100 |
| pooled est. (n=17) | Rga2<sub>(I)</sub> | 0.119 | 0.008 |
| pooled est. (n=12) | Rga2<sub>(II)</sub> | 0.278 | 0.042 |
| pooled est. (n=20) | GAP domain | 0.181 | 0.013 |

To investigate our hypothesis, we utilized Cdc24 mutants Cdc24-DH3 and Cdc24-DH5, which exhibit 2.5× and 10× reduction in their oligomerization capacity [Mionnet et al., 2008]. If the non-linear rate increase by Cdc24 (and thus its GEF activity) is linked to its oligomerization, we expect the oligomerization-reduced mutants to (1) show a reduced GEF activity and (2) boost the overall GTP hydrolysis rate *K* in a more linear fashion than Cdc24. A detailed discussion of our analysis is given in S2. In brief, Cdc24-DH5 did not express as full-length protein (S2 Fig. 1a). Cdc24-DH3 exhibited a 17× reduced GEF activity (Fig. 3c,e, Tab. 2) and increased the overall GTPase cycling rate *K* in a 1.7× more linear fashion than wildtype Cdc24 (S2 Tab. 1), confirming our hypothesis. Further, our data shows that previous findings on the *in vitro* GEF activity of peptides containing Cdc24 fragments do not translate to full-length Cdc24: Mionnet *et al*. found that a chemically induced oligomerization of peptides based on Cdc24 fragments does not affect their GEF activity, suggesting that the GEF activity might be oligomerization-independent [Mionnet et al., 2008]. We found that Cdc24-DH3, a Cdc24 mutant with reduced oligomerization capacity, has a reduced GEF activity, questioning the generalizability of data based on protein fragments.

In summary, the data show that Cdc24 is active at sub-µM concentrations, boosting the overall GTP hydrolysis rate in a non-linear fashion. We believe the non-linearity is linked to Cdc24 oligomerization and the subsequent release of Cdc24’s self-inhibition upon oligomerization.

### The GAP Rga2 self-inhibits upon oligomerization

Next to GEFs, GAPs also increase Cdc42 GTPase cycling, but through catalyzing the GTP hydrolysis step. We investigated the effect of the GAP Rga2, which so-far not yet been characterized as full-length protein. Rga2 has been shown to self-interact *in vivo* [Tarassov et al., 2008, Schlecht et al., 2012], but it is unclear if this interaction is direct or indirect (i.e. mediated via other proteins). Similar to Cdc24, Rga2 dimers or oligomers could affect its catalytic activity.

We first investigated Rga2 oligomerization using SEC-MALS experiments (Fig. 4a). Here a protein sample is separated by size under flow using size exclusion chromatography, where larger protein complexes elute first (at smaller volumes). The molecular weight of each fraction is then determined using multi-angle light scattering. We performed experiments using His-Rga2 (Rga2 that tagged with an N-terminal 6His purification tag, in the following referred to as Rga2_(*I*)_) and a peptide containing only the GAP domain of Rga2 [Smith et al., 2002]. In these *in vitro* experiments (Fig. 4a), Rga2_(*I*)_ was only detected as higher-order oligomers, with no monomers present. The GAP domain appeared as four closely overlapping peaks, suggesting the existence of higher than monomeric structures. Because the peaks overlapped so closely, the exact masses and thus oligomeric states could not be determined. The SEC-MALS data shows that Rga2 self-interacts directly, forming oligomers. Further, it implies that Rga2 oligomerization is facilitated by its C-terminal GAP domain.

**Figure 4.**
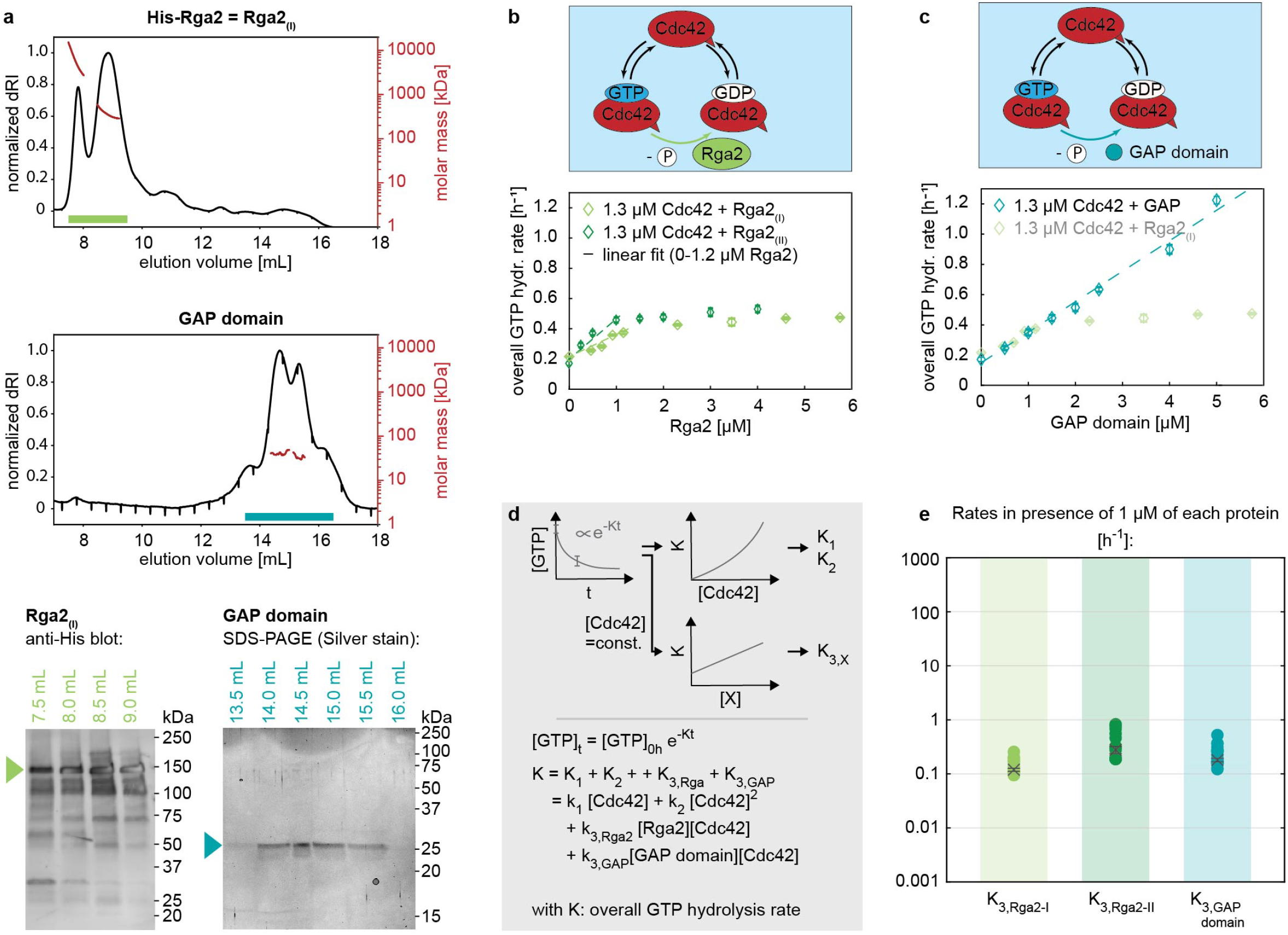
The saturation of the GAP activity of Rga2 may be linked to Rga2 oligomerization. (a) Size-exclusion profile and MALS analysis (top) and SDS-PAGE/ Western blot analysis of SEC-MALS elution fractions (bottom) of His-Rga2 (Rga2_(*I*)_) and Rga2’s GAP domain: Rga2 self-interacts, forming higher-order oligomeric structures. Rga2 oligomerization is facilitated by its C-terminal GAP domain, which in SEC-MALS elutes in multiple states (mixtures of monomers and higher than monomer structures). (b) Rga2_(*I*)_ and Rga2_(*II*)_ increase Cdc42 GTPase cycling in a linear fashion up until *∼*1.2 µM, after which their effect saturates. (c) Cdc42 GTPase cycling is accelerated by the GAP domain at a rate akin to Rga2, yet without its effect saturating. (d) Illustration of the data processing and fitting model. (e) Summary of the rates *K*: Rga2_(*I*)_, Rga2_(*II*)_ and the GAP domain have a similar rate. The values shown refer to the rate values in presence of 1 µM of each protein, e.g. ‘*K*_3,*Rga*2_’ refers to ‘*k*_3,*Rga*2_[Rga2][Cdc42]’ with [Cdc42]=[Rga2]=1 µM. Crosses with error bars represent the weighted mean and the standard error of the mean (S1), filled circles show individual measurements.

Next, we assessed the GAP activity of Rga2 through GTPase assays, examining Rga2’s concentration-dependent behavior. Rga2_(*I*)_ increases the overall GTP hydrolysis cycling rate *K* of Cdc42 in a linear fashion up until *∼*1.2 µM (corresponding to a 1:1 Rga2:Cdc42 ratio), after which its effect saturates (Fig. 4b). We also assessed a second Rga2 construct (His-Rga2-Flag, in the following referred to as Rga2_(*II*)_), which showed exactly the same behavior (Fig. 4b). One possible explanation is that at *∼*1.2 µM Rga2 the GTP hydrolysis step becomes rate-limiting; at this Rga2 concentration the hydrolysis reaches its maximum rate and cannot be further accelerated. Alternatively, at higher concentrations, Rga2 may self-inhibit. To investigate this, we analyzed the activity of Rga2’s GAP domain (Fig. 4c). The GAP domain displayed a similar linear rate increase, with rates approximately matching those of Rga2_(*I*)_ and Rga2_(*II*)_ (Fig. 4e, Tab. 2), yet without saturation, showing a linear increase across the full concentration range tested (up to 5 µM). This shows that the GTP hydrolysis step does not become rate-limiting at *∼*1 µM Rga2, supporting the possibility of Rga2 self-inhibition.

We speculate that the self-inhibition may originate from Rga2 oligomerization. Taken together, our data show that Rga2 has unique features distinct from the GAP domain, emphasizing that its full functionality is not confined to the GAP domain alone: (1) Rga2 self-interacts and forms oligomeric structures, facilitated by its C-terminal GAP domain. (2) The GAP activity of Rga2, unlike the activity of the GAP domain alone, saturates in GTPase assays. This saturation may be caused by self-inhibition upon oligomerization.

### Cdc24 and Rga2 synergistically regulate Cdc42 GTPase cycling

After having analyzed the individual effects of the GEF Cdc24 and the GAP Rga2, we explored their joint impact on Cdc42 GTPase cycling. This is noteworthy because (1) Cdc24 and Rga2 interact *in vivo* (directly or indirectly) [McCusker et al., 2007, Breitkreutz et al., 2010, Chollet et al., 2020] and (2) the interplay between GEFs and GAPs is theorized to play a significant role in G-protein signaling [Ross, 2008].

We conducted GTPase assays containing Cdc42 - Cdc24 - Rga2_(*I*)_ mixtures and fitted the data (Eq. 1, Fig. 5e) to obtain rates *K*. For such three-protein mixtures, we obtain *K*_3,*Cdc*24_ and *K*_3,*Rga*2_, which represent the individual rate contributions of the GEF and GAP, and the interaction term *K*_3,*Cdc*24,*Rga*2_. A positive interaction term represents synergy between the GEF and GAP, a negative value represents inhibition. If the term is zero, both proteins would not affect each other.

**Figure 5.**
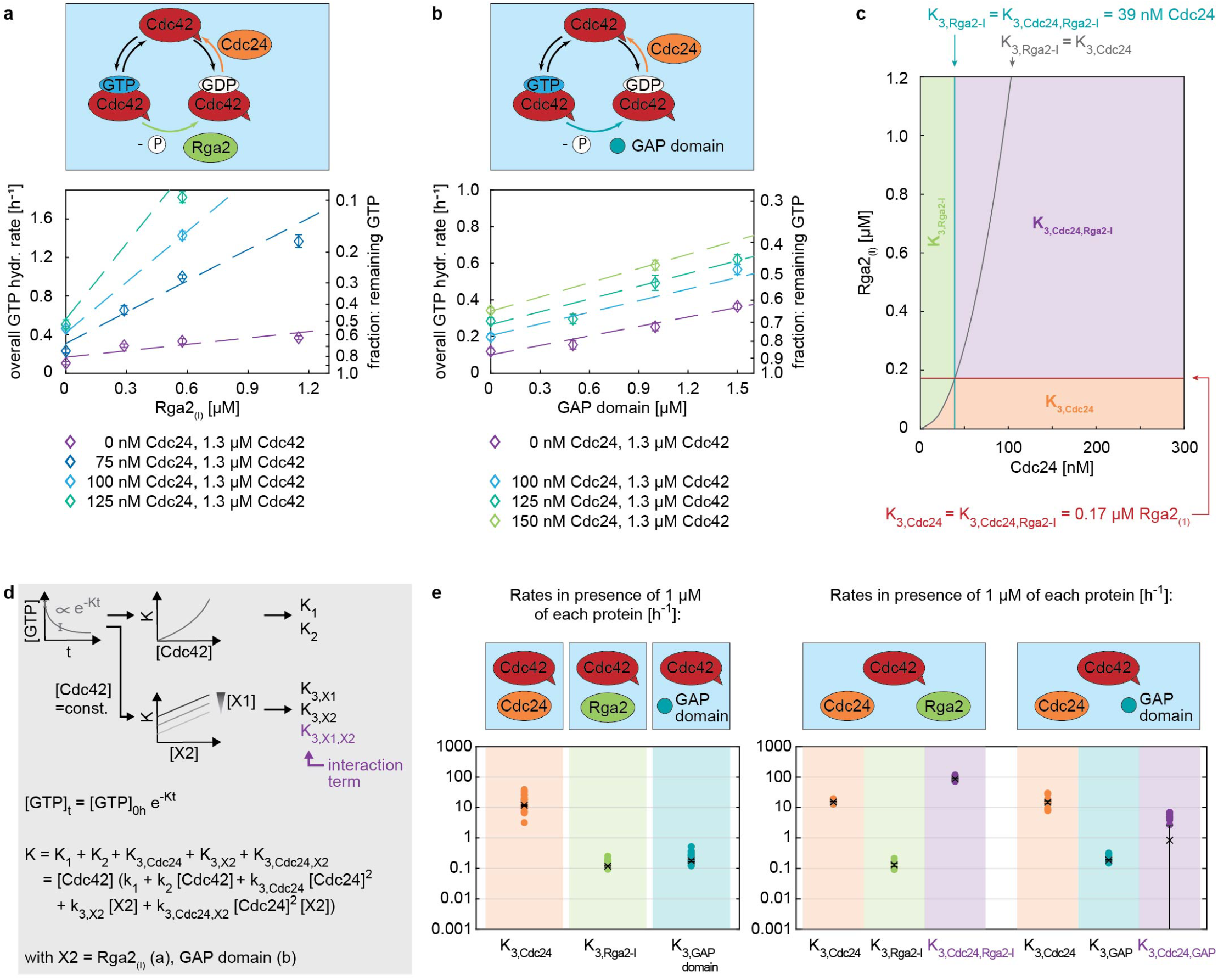
The GEF Cdc24 and GAP Rga2 regulate Cdc42 GTPase cycling synergistically. (a,b) GTPase assay data of (a) Cdc42-Cdc24-Rga2_(*I*)_ and (b) Cdc42-Cdc24-GAP domain mixtures. (c) Regime diagram illustrating the dominant rate *K* across varying Cdc24 and Rga2_(*I*)_ concentrations; above 39 nM Cdc24 and 0.17 µM Rga2, the interaction term *K*_3,*Cdc*24,*Rga*2_ becomes predominant. The diagram was generated using rates *k*_3_ for effectors Cdc24 and Rga2_(*I*)_ given in Tab. 3. An extended diagram is shown in S6. (d) Illustration of the data processing and fitting model. (e) Summary of the rates obtained in the three-protein assay (right) in comparison to those of the two-protein assay (left): In three-protein assays the rate contribution of the individual proteins is comparable to those obtained in the two-protein assay. Additionally, an interaction rate is obtained (shown in purple). This rate is large in the case of Cdc42-Cdc24-Rga2_(*I*)_ mixtures (*K*_3,*Cdc*24,*Rga*2_ ≈ 87), indicating a synergy between Cdc24 and Rga2_(*I*)_. For Cdc42-Cdc24-GAP domain mixtures the interaction rate is zero (*K*_3,*Cdc*24,*GAP domain*_ ≈ 0), revealing no synergy. The values shown refer to the rate values in presence of 1 µM of each protein, e.g. ‘*K*_3,*Cdc*24_’ refers to ‘*k*_3,*Cdc*24_ [Cdc24]^2^ [Cdc42]’ with [Cdc42]=[Cdc24]=1 µM. Crosses with error bars represent the weighted mean and the standard error of the mean (S1), and filled circles show individual measurements.

In presence of both Cdc24 and Rga2_(*I*)_, the GTPase activity of Cdc42 increases drastically (Fig. 5a). The contributions of the individual proteins (*K*_3,*Cdc*24_, *K*_3,*Rga*2_) are roughly the same as in assays containing only one of the effectors (Fig. 5d, Tab. 3). Importantly, the interaction term *K*_3,*Cdc*24,*Rga*2_ is the dominating term for essentially the entire concentration regime (i.e. above 0.17 µM Rga2_(*I*)_ and 39 nM Cdc24, Fig. 5c), suggesting a GEF - GAP synergy.

**Table 3.** Cdc42 - effector protein X interaction rates 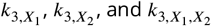.

| | Effector protein $X_1$ | Effector protein $X_2$ | $k_{3,X_1}$<br>[$\mu M^{-3} h^{-1}$] | $k_{3,X_1}$<br>std. err. | $k_{3,X_2}$<br>[$\mu M^{-2} h^{-1}$] | $k_{3,X_2}$<br>std. err. | $k_{3,X_1,X_2}$<br>[$\mu M^{-4} h^{-1}$] | $k_{3,X_1,X_2}$<br>std. err. |
| --- | --- | --- | --- | --- | --- | --- | --- | --- |
| pooled est. (n=7) | Cdc24 | Rga2<sub>(I)</sub> | 15.126 | 0.736 | 0.131 | 0.015 | 86.961 | 6.055 |
| pooled est. (n=7) | Cdc24 | Rga2<sub>(II)</sub> | 13.001 | 2.961 | 0.481 | 0.090 | 51.702 | 4.827 |
| pooled est. (n=11) | Cdc24 | GAP domain | 14.895 | 1.579 | 0.188 | 0.012 | 0.842 | 1.674 |

To ensure that a positive *K*_3,*Cdc*24,*Rga*2_ is not an assay artifact, we performed additional GTPase assays using proteins considered inert (bovine serum albumin (BSA), Casein), as well as mixtures of BSA with Cdc24 and BSA with Rga2_(*I*)_ (S4). In these assays no significant synergy occurred, confirming that *K*_3,*Cdc*24,*Rga*2_ is a genuine signal.

The positive *K*_3,*Cdc*24,*Rga*2_ could originate from two sources: (1) Cdc24-Rga2 synergy due to rate-limiting effects of different steps in the GTPase cycle, and (2) Cdc24-Rga2 synergy due to protein-protein interactions.

Could rate-limiting steps be the *only* source of the positive *K*_3,*Cdc*24,*Rga*2_? The rate-limiting step model is a common framework used to describe the cycling behavior of GTPases. It assumes that at least one of the three GTPase cycle steps — GTP binding, GTP hydrolysis, or GDP release — is rate-limiting. It has the potential to explain the synergy term we observed in our data: if both the GDP release step, accelerated by the GEF Cdc24, and the GTP hydrolysis step, accelerated by the GAP Rga2, are rate-limiting, then adding both effectors would accelerate the overall cycle more than either alone. This synergistic increase in cycling speed would result from their combined effect on the GTPase cycle and not from direct interaction between the proteins. The effect size of the synergetic term, as predicted by the rate-limiting step model, depends on the concentrations of the GEF and GAP used: the synergy is strongest when both steps — GDP release and GTP hydrolysis — are rate-limiting, which occurs at higher concentrations of GEF and GAP (Fig. 6b). Conversely, the synergy diminishes when these steps are not rate-limiting, i.e. at lower effector concentrations (Fig. 6a). At which GEF and GAP concentrations do GDP release and GTP hydrolysis become rate limiting? When either step becomes rate-limiting, we expect the overall GTP hydrolysis rate *K* to saturate (Fig. 6c). We conducted experiments with only one effector present (Cdc42 + Cdc24, Cdc42 + Rga2): we do not observe saturation of *K* with increasing Cdc24 concentration (Fig. 3b), suggesting that GDP release does not become rate-limiting within our tested concentration range. Although *K* does saturate with full-length Rga2 (Fig. 4b), it does not with the GAP domain (Fig. 4c), suggesting that we also do not reach concentration regimes where GTP hydrolysis becomes rate-limiting. Thus, we expect the overall synergy due to rate-limiting steps to be small. Additionally, in the rate-limiting step model the maximal possible accelerations by the GEF and GAP are interconnected, i.e. the maximal acceleration through the GEF limits the maximal acceleration that can be achieved by the GAP (and vice versa). We analyzed if the accelerations that we observe by the GEF and GAP fit this model (S5). In brief, we experimentally observe significantly larger accelerations by the GEF and GAP alone than the rate limiting step model allows for, also suggesting that it cannot fully explain the synergy term in our data.

**Figure 6.**
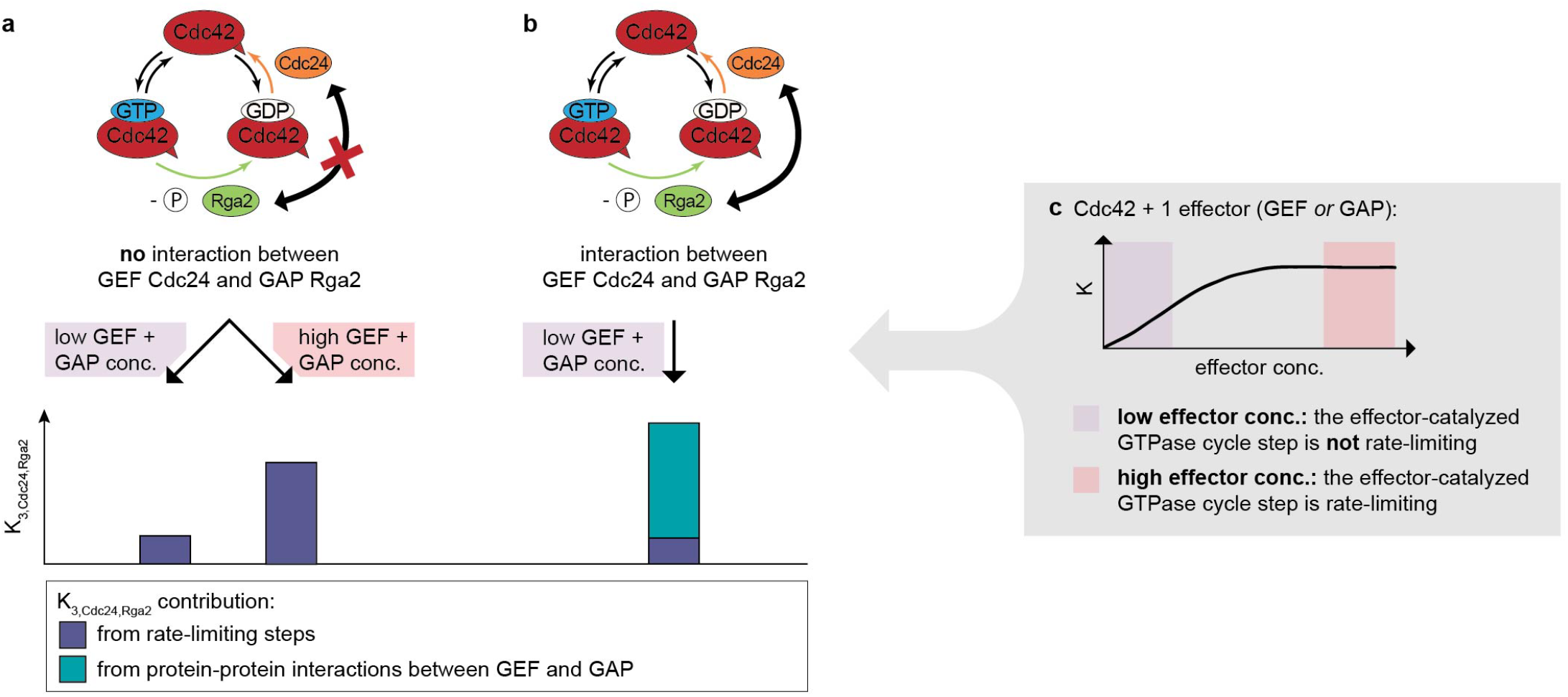
GEF-GAP synergies can originate from rate-limiting steps and specific GEF-GAP interactions. (a) In the absence of an GEF-GAP interaction, synergy arises solely from rate-limiting steps. At low effector concentrations the GEF-and GAP-catalyzed GTPase cycle steps are not in the rate-limiting regime (see (c)), resulting in a small synergy. At high effector concentrations the GEF-and GAP-catalyzed GTPase cycle steps are in the rate-limiting regime, resulting in a larger synergy (see (c)). (b) In presence of a specific GEF-GAP-interaction, synergy can additionally arise from protein-protein interactions. At low effector concentrations the contribution of the synergy from protein-protein interactions dominates, as the contribution from rate-limiting steps in this concentration regime is low. (c) GEF and GAP concentrations define rate-limiting step regimes: at low effector concentrations (purple regime), the effector-catalyzed step is not rate-limiting, and increases in effector concentrations lead to a higher overall GTPase hydrolysis rate *K*. At high effector concentrations (red regime), the effector the effector-catalyzed step is rate-limiting, thereby capping the overall GTPase hydrolysis rate *K*.

More importantly, we carried out GTPase assays with Cdc24 and the GAP domain (Fig. 5b) and observed that here the interaction term *K*_3,*Cdc*24,*GAP*_ is close to zero (Fig. 5d, Tab. 3). If the rate-limiting step model would fully account for the synergy term in our data, *K*_3,*Cdc*24,*GAP*_ would be expected to be close to *K*_3,*Cdc*24,*Rga*2_, given the comparable rates of the GAP domain and Rga2_(*I*)_. This strongly supports that *K*_3,*Cdc*24,*Rga*2_ cannot be explained by the rate-limiting step model alone and points to a protein-specific synergy between Cdc24 and Rga2. We suspect that *K*_3,*Cdc*24,*GAP*_ is close to, but not exactly, zero, due to synergistic effects arising from rate-limiting steps in the GTPase cycle. However, because the concentrations of Cdc24 and GAP used are well outside the regimes where GDP release or GTP hydrolysis become rate-limiting, these effects contribute minimally to the overall synergy, resulting in a *K*_3,*Cdc*24,*GAP*_ close to zero.

Taken together, we observe a positive interaction term *K*_3,*Cdc*24,*Rga*2_ in our data, which cannot be fully explained by rate-limiting steps in the GTPase cycle alone and instead points to a protein-specific synergy between Cdc24 and Rga2. Next, we used SEC-MALS to investigate Cdc24-Rga2_(*I*)_ binding as a potential origin of the synergy (Fig. 7): We did not observe Cdc24-Rga2 heterodimers, as the Rga2 oligomer peak lacked Cdc24 (Fig. 7, Western blot). However, the Cdc24 peak shifted towards a higher molecular weight, suggesting that an Rga2 fragment binds to Cdc24. This implies weak binding, as no Cdc24-Rga2 dimers or oligomers survived the flow under which SEC-MALS is performed. We analyzed the SEC-MALS fractions using several SDS-PAGE staining techniques and Western blotting. We detected a *∼*25 kDa fragment of Rga2 in the Cdc24 peak, visible only with SYPRO Ruby staining (Fig. 7, SYPRO Ruby (blue arrow)), revealing it lacks both the N-terminal His-tag (detected via anti-His Western blotting) and tryptophan residues (detected via stain-free SDS-PAGE). We analyzed the Rga2 sequence to determine if the lack of tryptophan could indicate which Rga2 domain binds to Cdc24, but all tryptophan-devoid Rga2 fragments are at least 25 kDa, meaning any of them could match the fragment observed on gel (S7). Therefore, the binding region remains unclear. We also conducted GTPase assays with Cdc24 and Rga2_(*II*)_, which again showed synergy, although 40% reduced (S8 and Tab. 3). We suspect that the C-terminal Flag-tag of Rga2_(*II*)_, which is not present in Rga2_(*I*)_, may weaken Cdc24-Rga2 binding, resulting in a reduced synergy between the proteins. This would imply that Rga2’s C-terminus is involved in Cdc24-Rga2 binding.

**Figure 7.**
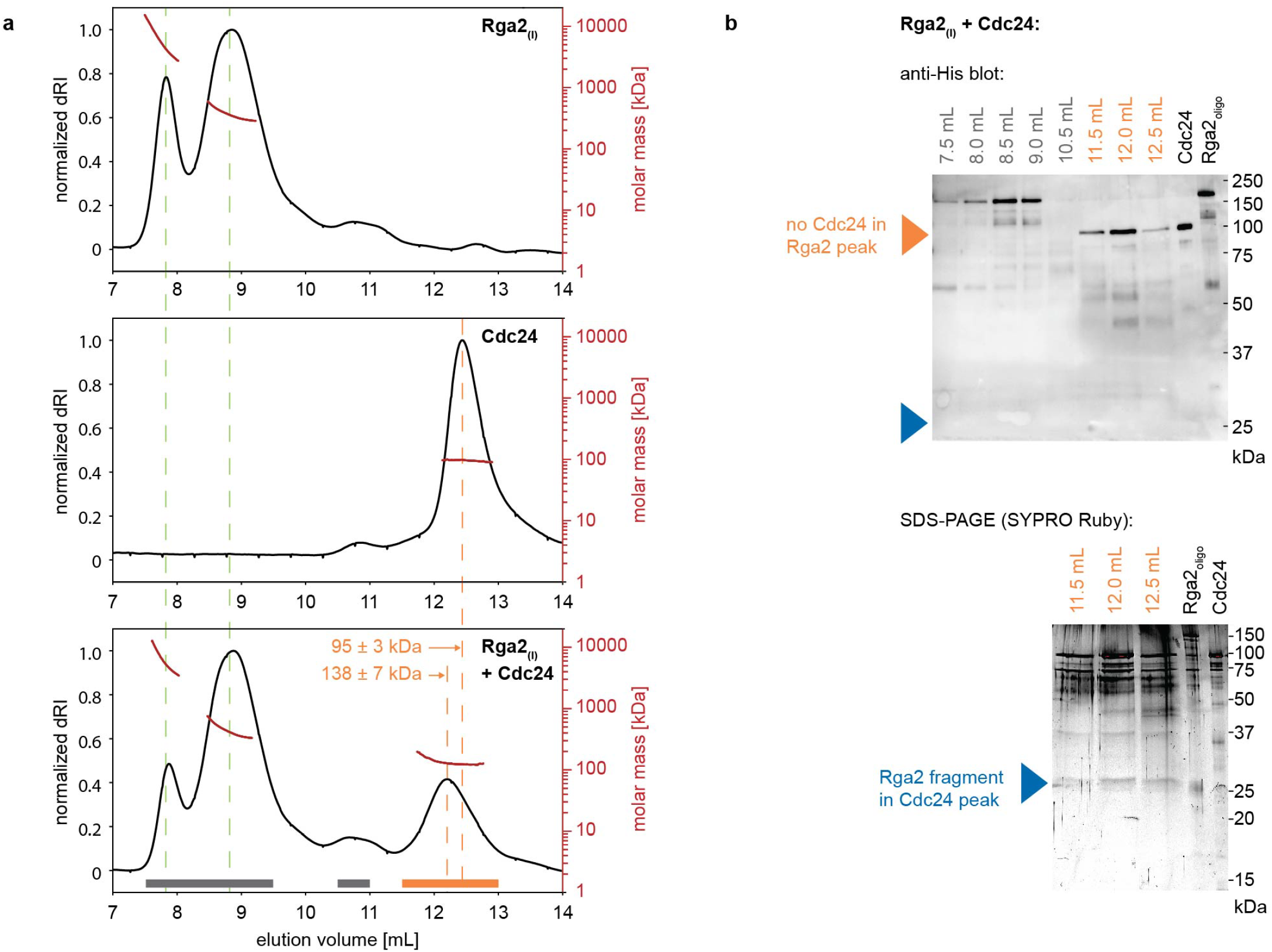
The GEF Cdc24 and GAP Rga2 bind through weak interactions. Size-exclusion profile and MALS analysis (a) and SDS-PAGE/ Western blot analysis of SEC-MALS elution fractions (b) of Rga2_(*I*)_, Cdc24, and Rga2_(*I*)_ - Cdc24 mixtures: In SEC-MALS experiments with Cdc24, Rga2_(*I*)_, and their mixtures, full-length Cdc24 is not present in Rga2 peaks (Western blot: orange arrow). Instead, binding occurs with an Rga2_(*I*)_ fragment (*∼*25 kDa, SDS-PAGE: blue arrow), which shifts the Cdc24 peak towards larger molecular weights. This Rga2 fragment is devoid of tryptophan and lacks a His-tag (Western blot: blue arrow).

In conclusion, our data reveals a GEF-GAP synergy that is specific to Rga2 and Cdc24, absent in assays containing Cdc24 and the GAP domain. This synergy appears to arise from weak Cdc24-Rga2 binding, through which Cdc24 might alleviate the self-inhibition of oligomeric Rga2 (Fig. 8).

**Figure 8.**
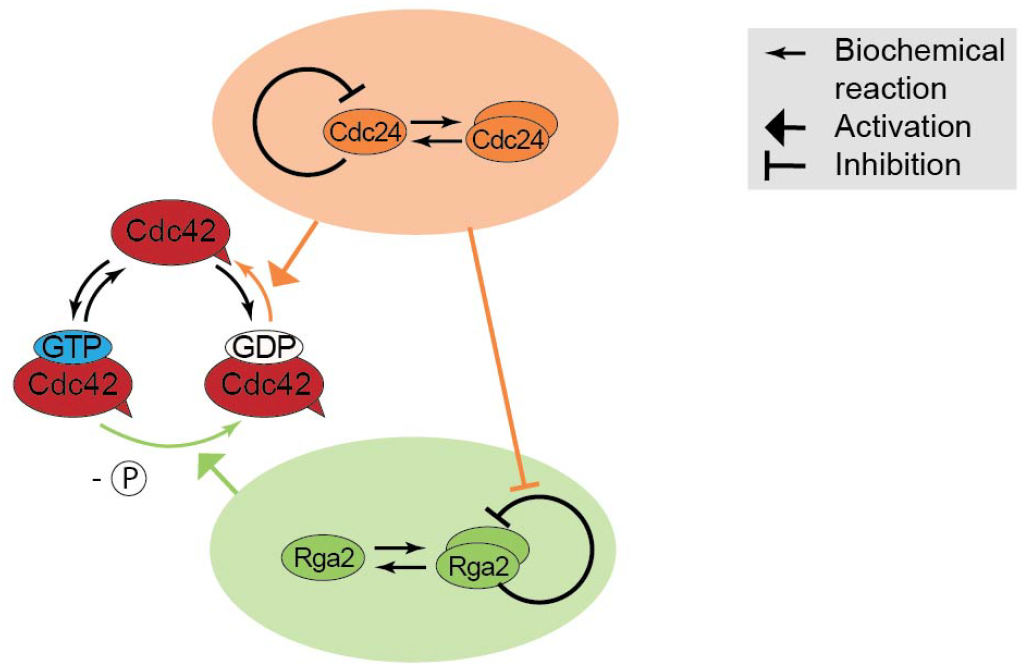
Oligomerization-driven regulation of Cdc42 GTPase activity. Speculative model of Cdc42 GTPase activity regulation through the GEF Cdc24 and GAP Rga2: Both Cdc24 and Rga2 form oligomers. Cdc24’s self-inhibition is released upon oligomerization, increasing its GEF activity. In contrast, Rga2 self-inhibits upon oligomerization. Together, Cdc24 and Rga2 synergize, due to Cdc24 alleviating the self-inhibition of oligomeric Rga2.

## Discussion

Synergy in GTPase regulation can arise through distinct mechanisms. One synergy arises from the acceleration of rate-limiting steps in the GTPase cycle by GEFs and GAPs: when both GDP release and GTP hydrolysis are rate-limiting, the combined action of GEFs and GAPs can enhance the cycling rate more than either effector alone. Our data points to an additional form of synergy, which is based on protein-protein interactions: we observed a synergistic effect between Cdc24 and Rga2, which is absent in data where Cdc24 is combined with the isolated GAP domain. Our data suggests a novel functional coupling between GEFs and GAPs that arises due to protein-specific interactions between Cdc24 and Rga2. This underscores the importance of considering the molecular context when dissecting GTPase regulatory networks.

Our findings indicate that oligomerization is a critical factor shaping the GEF and GAP activities of Cdc24 and Rga2 (Fig. 8a): Cdc24’s GEF activity increases non-linearly with concentration, and data from an oligomerization-reduced mutant suggest that this non-linearity may result from the release of self-inhibition upon oligomerization [Shimada et al., 2004], similar to the activity-enhancing effect of Bem1 [Rapali et al., 2017]. Rga2 appears also to self-inhibit, as indicated by the saturation of its activity observed in the data. Interestingly, Rga2’s self-inhibition might originate from oligomerization and be alleviated by Cdc24. We suspect that this is the source of the GEF-GAP synergy, consistent with our finding that Cdc24 and the GAP domain lack synergy, as the GAP domain’s activity does not saturate (i.e. does not self-inhibit). Taken together, our data indicates that oligomerization has opposing effects on the GEF Cdc24 and the GAP Rga2; while Cdc24 overcomes self-inhibition, Rga2 self-inhibits. Subsequently Rga2 relies on Cdc24 to overcome its self-inhibition, suggesting a functional connection between GAP activity and the GEF.

The emergence of Rga2’s dependence on Cdc24 to release its self-inhibition could be the result of epistasis-a phenomenon where the effect of one gene masks or modifies the effect of another gene: Cdc24 is an essential gene product in *S. cerevisiae* and highly conserved in the fungal tree [Diepeveen et al., 2018], making it a persistent player in the polarity network. Mutations in Rga2 that result in self-inhibition would be permitted and masked by Rga2’s synergistic effect with Cdc24, and could emerge without reducing the organism’s fitness. Rga2 is only one of four GAPs of Cdc42 (among Rga1, Bem2, and Bem3). It was suggested that each GAP plays a distinct role in Cdc42 regulation, of which the level of GAP activity could be a part of [Smith et al., 2002]. The role of GAP oligomerization, self-inhibition, and interaction with the GEF and other regulatory factors could be other distinguishing factors. *In vitro* analysis of other GAPs would help to clarify if and how these factors contribute to different roles of GAPs in *S. cerevisiae*.

Our data exemplifies non-linearities of Cdc42 GTPase cycle regulation: (1) the overall GTP hydrolysis rate of Cdc42 increases quadratically with Cdc24 concentration, and (2) the GEF Cdc24 and GAP Rga2 exhibit synergy. Both non-linearities could contribute to establishing polarity through creating regimes of high and low Cdc42 activity: (1) Temporal regulation of Cdc42 activity: *In vivo*, the timed release of Cdc24 from the nucleus (thus suddenly increasing the effective Cdc24 concentration) is known to be part of the polarity trigger [Shimada et al., 2000]. We suspect that the non-linear increase of the overall GTP hydrolysis rate of Cdc42 with Cdc24 concentration is a mechanistic element of Cdc24’s function *in vivo*: once Cdc24 gets released from the nucleus, the GTPase cycling speed of Cdc42 increases strongly and suddenly (due to its non-linear dependence on Cdc24 concentration). If Cdc24 additionally releases Rga2’s self-inhibition and increases its GAP activity, Cdc24 further boosts Cdc42 GTPase cycling through activating Rga2. Timed by the release of Cdc24 from the nucleus, cells can quickly transition from a regime of low GTPase activity of Cdc42 (before the release of Cdc24) to a regime of high GTPase activity of Cdc42 (after the release of Cdc24). This sudden change in Cdc42’s GTPase cycling speed could be part of the trigger that initiates polarity establishment. (2) Spatial regulation of Cdc42 activity: The synergistic regulation of Cdc42’s GTPase activity through GEFs and GAPs a resourceful and advantageous way of regulation; if regulatory factors have a synergistic interplay, wide ranges of up-regulation can be achieved through a small number of components. This synergy also implies that Cdc42 has a significantly higher GTPase activity at the polarity spot, where it is surrounded by many effector proteins, that also regulate each other. We suspect the strong up-regulation at the site of bud emergence and the rather low baseline activity at other sites to have a cellular purpose, and imagine it is contributing to Cdc42 accumulation.

Beyond its impact on our understanding of the yeast system, our findings may also apply to other eukaryotes: Cdc42 is highly conserved among eukaryotes, and plays a central role in polarity establishment in many of these [Nelson, 2003, Etienne-Manneville, 2004, Thompson, 2013, Diepeveen et al., 2018]. We imagine that the principles of its regulation are also conserved there, and propose that synergies between GEFs and GAPs also occur in these systems.

## Materials and Methods

### Plasmid construction

Genes of interest (Cdc42, Rga2) were obtained from the genome of *Saccharomyces cerevisiae* W303 and were amplified through PCR. The target vector (pET28a-His-mcm10-Sortase-Flag, received from N. Dekker (TU Delft) and based on pBP6 [Douglas and Diffley, 2016]) was also amplified through PCR. Additionally, each PCR incorporated small homologous sequences needed for Gibson assembly [Gibson et al., 2009]. After Gibson assembly, the resulting mixture was used to transform chemically competent Dh5*α* and BL21 DE::3 pLysS cells and plated out onto a Petri dish containing Lysogeny broth agar and the correct antibiotic marker.

Construction routes of plasmids and primers used for each PCR are shown in S9. The amino acid sequences of the proteins used in this publication are stated in S10 and their design is discussed in detail in [Tschirpke et al., 2023b].

### Buffer composition

If not mentioned otherwise, buffers are of the composition stated in Tab. 5.

**Table 4.** List of protein constructs/plasmids used throughout this publication.

| Plasmid | Description | Source |
| --- | --- | --- |
| pRV007 | Cdc42 | this work |
| pRV014 | Rga2<sub>(I)</sub> | this work |
| pRV110 | Rga2<sub>(II)</sub> | this work |
| pRV138 | GAP domain (aa 797-981 from Rga2 [Smith et al., 2002]) | this work |
| pDM272 | Cdc24 | received from D. McCusker (University of Bordeaux) [Rapali et al., 2017] |
| pRV136 | Cdc24-DH5 (incl. mutation F322A [Mionnet et al., 2008]) | this work |
| pRV137 | Cdc24-DH3 (incl. mutation L339A, E340A [Mionnet et al., 2008]) | this work |

**Table 5.** Buffer composition.

| Buffer | Composition |
| --- | --- |
| Lysis buffer | 50 mM Tris-HCl (pH=8.0), 1 M NaCl, 5 mM imidazole, 1 mM 2-mercaptoethanol, supplemented with EDTA-free Protease inhibitor cocktail (Roche) and 1 mM freshly prepared PMSF. |
| His-AC washing buffer | 50 mM Tris-HCl (pH=8.0), 1 M NaCl, 5 mM imidazole, 1 mM 2-mercaptoethanol. |
| His-AC elution buffer | 50 mM Tris-HCl (pH=8.0), 100 mM NaCl, 500 mM imidazole, 1 mM 2-mercaptoethanol. |
| SEC buffer | 50 mM Tris-HCl (pH=7.5), 100 mM NaCl, 10 mM MgCl<sub>2</sub>, 1 mM 2-mercaptoethanol. |

### Protein expression and purification

**Cdc42 (pRV007)** was expressed in Bl21::DE3 pLysS cells. Cells were grown in Lysogeny broth at 37^°^C until an OD_600_ of 0.7. The expression was induced through addition of 1.0 mM IPTG, after which cells were grown for 3 h at 37^°^C. Cells were harvested through centrifugation. Cell pellets were resuspended in lysis buffer and lysed with a high-pressure homogenizer (French press cell disruptor, CF1 series Constant Systems) at 4^°^C, using 5-10 rounds of exposing the sample to pressurization. The cell lysate was centrifuged at 37000× g for 30 min and the supernatant was loaded onto a HisTrap™ excel column (Cytiva).

After several rounds of washing with His-AC washing buffer, the protein was eluted in a gradient of His-AC washing buffer and His-AC elution buffer. The protein as dialyzed twice in SEC buffer. After the addition of 10% glycerol, samples were flash frozen in liquid nitrogen and kept at-80^°^C for storage.

The expression and purification of **Cdc24 (pDM272)** is, with modifications, based on the protocol described previously [Rapali et al., 2017]: Cdc24 was expressed in Bl21::DE3 pLysS cells. Cells were grown in Lysogeny broth at 37^°^C until an OD_600_ of 0.7. The expression was induced through addition of 0.2 mM IPTG, after which cells were grown for 18 h at 18^°^C. Cells were harvested through centrifugation. Cell pellets were resuspended in lysis buffer and lysed with a high-pressure homogenizer (French press cell disruptor, CF1 series Constant Systems) at 4^°^C, using 5-10 rounds of exposing the sample to pressurization. The cell lysate was centrifuged at 37000× g for 30 min and the supernatant was loaded onto a HisTrap™ excel column (Cytiva). After several rounds of washing with His-AC washing buffer, the protein was eluted in a gradient of His-AC washing buffer and His-AC elution buffer. The sample was further purified by size exclusion chromatography using SEC buffer and a HiPrep 16/60 Sephacryl S-300 HR (Cytiva) column. Fractions containing full-size protein were concentrated using Amicon®Ultra 4 mL centrifugal filters (Merck). After the addition of 10% glycerol, samples were flash frozen in liquid nitrogen and kept at-80^°^C for storage.

Cdc24 mutants **Cdc24-DH3 (pRV137)** and **Cdc24-DH5 (pRV136)** were expressed and purified in the same fashion as Cdc24 (pDM272), with the only modification that Lysis buffer, His-AC washing buffer, and His-AC elution buffer were all supplemented with 0.1% Tween-20, 0.1% NP-40, and 0.1% Triton-X-100.

**Rga2**_(*I*)_ **(pRV014)** and **Rga2**_(*II*)_ **(pRV110)** were expressed in Bl21::DE3 pLysS cells. Cells were grown in Lysogeny broth at 37^°^C until an OD_600_ of 0.7. The expression was induced through addition of 0.2 mM IPTG, after which cells were grown for 18-24 h at 10^°^C. Cells were harvested through centrifugation. Cell pellets were resuspended in lysis buffer (supplemented with 0.1% Tween-20, 0.1% NP-40, and 0.1% Triton-X-100) and lysed with a high-pressure homogenizer (French press cell disruptor, CF1 series Constant Systems) at 4^°^C, using 5-10 rounds of exposing the sample to pressurization. The cell lysate was centrifuged at 37000× g for 30 min and the supernatant was loaded onto a HisTrap™ excel column (Cytiva). After several rounds of washing with His-AC washing buffer (supplemented with 0.1% Tween-20, 0.1% NP-40, and 0.1% Triton-X-100), the protein was eluted in a gradient of His-AC washing buffer and His-AC elution buffer (both supplemented with 0.1% Tween-20, 0.1% NP-40, and 0.1% Triton-X-100). The sample was further purified by size exclusion chromatography using SEC buffer and a HiPrep 16/60 Sephacryl S-300 HR (Cytiva) column. Fractions containing full-size protein were concentrated using Amicon®Ultra 4 mL centrifugal filters (Merck). After the addition of 10% glycerol, samples were flash frozen in liquid nitrogen and kept at-80^°^C for storage.

**GAP domain (pRV138)** was expressed in Bl21::DE3 pLysS cells. Cells were grown in Lysogeny broth at 37^°^C until an OD_600_ of 0.7. The expression was induced through addition of 1.0 mM IPTG, after which cells were grown for 3 h at 37^°^C. Cells were harvested through centrifugation. Cell pellets were resuspended in lysis buffer and lysed with a high-pressure homogenizer (French press cell disruptor, CF1 series Constant Systems) at 4^°^C, using 5-10 rounds of exposing the sample to pressurization. The cell lysate was centrifuged at 37000× g for 30 min and the supernatant was loaded onto a HisTrap™ excel column (Cytiva). After several rounds of washing with His-AC washing buffer, the protein was eluted in a gradient of His-AC washing buffer and His-AC elution buffer. The sample was further purified by size exclusion chromatography using SEC buffer and a HiPrep 16/60 Sephacryl S-300 HR (Cytiva) column. Fractions containing full-size protein were concentrated using Amicon®Ultra 4 mL centrifugal filters (Merck). After the addition of 10% glycerol, samples were flash frozen in liquid nitrogen and kept at-80^°^C for storage.

**Casein, Bovine serum albumin (BSA), and Ras** were purchased. Casein (Sigma-Aldrich, cat. no. C7078) was dissolved in SEC buffer. BSA (23209, Thermo Scientific) was dialyzed twice in SEC buffer. Ras (human) (EMD Millipore, cat. no. 553325) was diluted in SEC buffer.

All proteins are shown on SDS-PAGE in S11. To determine **protein concentrations**, samples were treated with Compat-Able Protein Assay Preparation Reagent Kit (Thermo Fischer Scientific) and analyzed using a BCA assay (Pierce BCA Protein Assay Kit, Thermo Fischer Scientific).

### Note on His-affinity chromatography

HisTrap™ excel column (Cytiva) columns bought 2020 or later required a higher amount of imidazole in the lysis and washing buffer that stated in Tab. 5, as indicated by the recommendation ‘use 20-40 mM imidazole in sample and binding buffer for highest purity’ on the column package. For these columns the amount of imidazole in lysis and His-AC washing buffer was increased to 50 mM.

### GTPase assay

GTPase activity was measured using the GTPase-Glo™ assay (Promega), following the steps described in the assay manual and in [Tschirpke et al., 2024]. In brief, 5 µL protein in SEC buffer (Tab. 5) was mixed with 5 µL of a 2× GTP-solution (10 µM GTP, 50 mM Tris-HCl (pH=7.5), 100 mM NaCl, 10 mM MgCl_2_, 1 mM 2-mercaptoethanol (Sigma Aldrich), 1 mM dithiothreitol in 384-well plates (Corning) to initiate the reaction. The reaction mixture was incubated for 60 to 100 min at 30^°^C on an Innova 2300 platform shaker (New Brunswick Scientific) (120 rpm), before the addition of 10 µL Glo buffer and another 30 min incubation. The Glo buffer contains a nucleoside-diphosphate kinase that convert remaining GTP to ATP. Addition of 20 µL detection reagent, containing a luciferase/luciferin mixture, makes the ATP luminescent, which was read on a Synergy HTX plate reader (BioTek) in luminescence mode. Each data point shown corresponds to the average of 3-4 replica samples per assay.

This GTPase assay can be sensitive to small concentration differences, especially of effector proteins. To reduce the variability between assays and to increase comparability of different assay sets, 6× proteins stocks were made using serial dilutions (with SEC buffer). The assays were conducted using the same 6× proteins stocks within a few days, during which these stocks were kept at 4^°^C. For each assay, equivalent volumes of 6× protein stocks were diluted to 2× mixtures (e.g. 10 µL Cdc42 + 20 µL SEC buffer, 10 µL Cdc42 + 10 µL effector protein 1 + 10 µL SEC buffer, 10 µL Cdc42 + 10 µL effector protein 1 + 10 µL effector protein 2,…). Incubation of 5 µL of this protein mixture with 5 µL of 2× GTP solution, as described above, resulted in the concentrations stated in the figures.

We first assessed Cdc42 alone using several Cdc42 concentration. We then examined the individual Cdc42 – effector mixtures (two-protein assays), using one Cdc42 concentration and several effector concentrations. We then conducted assays using Cdc42 and two effectors. These three-protein assays contain wells with Cdc42, Cdc42 + effector 1, Cdc42 + effector 2, Cdc42 + effector 1 + effector 2, and additional ‘buffer’ wells used for normalization. For feasibility the three-protein assays include a reduced number of effector concentrations (in comparison to the two-protein assays).

### Fitting of GTPase assay data

The GTPase model, analysis procedure, and analysis code are described in detail in [Tschirpke et al., 2024] and available at [Tschirpke et al., 2023a]. Key aspects are summarized here:

The amount of remaining GTP correlates with the measured luminescence. Wells without protein (‘buffer’) were used for the normalization and represent 0% GTP hydrolysis:

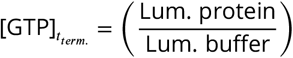

Wells where no GTP was added showed luminescence values corresponding to <1% remaining GTP. Given the small deviation to 0%, and that GTPase reactions of protein mixtures leading to <5% remaining GTP were excluded from further analysis [Tschirpke et al., 2024], we did not normalize the data using luminescence values corresponding to 0% GTP.

Reactions were carried out with three to four replicates (wells) per assay, and the average (‘Lum.’) and standard error of the mean (‘ΔLum.’) of each set was used to calculate the amount of remaining GTP at the time of reaction termination and the error of each set:

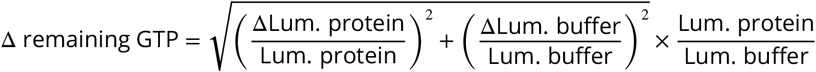

The data was fitted using a GTPase activity model (described in brief in S1 and in detail in [Tschirpke et al., 2024]) where the GTP decline occurring during the GTPase reaction is approximated with an exponential:

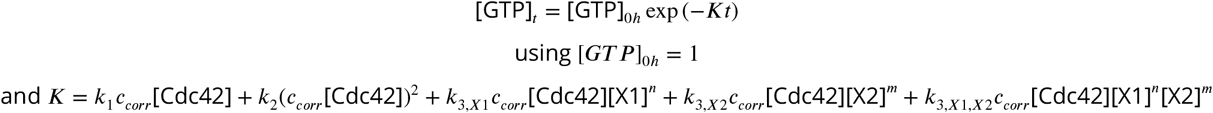

Here *K* represents the overall GTP hydrolysis rate, *X*1 and *X*2 are effector proteins with *n* and *m* = (1, 2}, and *c*_*corr*_ is a variable used to map all factors that lead to variations between GTPase assays onto the Cdc42 concentration. The pooled estimates of rates *k*_1_, *k*_2_, and *k*_3_ were determined through weighting their standard error.

### SEC-MALS

The oligomerization states of Cdc24, Rga2, the GAP domain, and mixtures thereof, were estimated using analytical size exclusion chromatography coupled to multi-angle light scattering (SEC-MALS). Purified protein samples were resolved on a Superdex 200 Increase 10/300 GL column (Cytiva) connected to a high-performance liquid chromatography (HPLC) unit (1260 Infinity II, Agilent) running in series with an online UV detector (1260 Infinity II VWD, Agilent), an 8-angle static light scattering detector (DAWN HELEOS 8+; Wyatt Technology), and a refractometer (Optilab T-rEX; Wyatt technology). Experiments were performed using SEC buffer supplemented with 0.02% natrium azide and filtered through 0.1 µM pore filters (Durapore Membrane Filter 0.1 µM, Millipore). Protein samples were prepared at a total volume of 60 µL and final concentrations of 5.7 - 7.1 µM. After mixing, samples were incubated on ice for 30 min and then spun down at 21000× g for 10 min (Eppendorf Centrifuge 5424 R). 50 µL of protein sample was injected onto the column and eluted using SEC buffer supplemented with 0.02% natrium azide at 0.5 mL/min. Peak fractions were collected. 1 mg/mL monomeric BSA (Albumin monomer bovine, Sigma-Aldrich) was used as a reference for mass calibration. All experiments were performed as triplicates.

### SDS-PAGE

SDS-PAGE gels (12-15% acrylamide) were prepared freshly. In brief, a solution of 375 mM Tris-HCl (pH=8.8), 30-37.5 v/v% 40% acrylamide solution (Biorad), 0.2 w/v% sodium dodecyl sulfate (Sigma Aldrich), 0.5 v/v% 2,2,2-Trichloroethanol (Sigma Aldrich), 0.1 w/v% ammonium persulfate (Sigma Aldrich), and 0.1 v/v% N,N,N’,N’-tetramethyl ethylenediamine (Sigma Aldrich) was prepared and casted into 1.00 mm mini-protean glass plates (Biorad), filling them up to 80%. To protect the gel surface from drying, a layer of isopropanol (Sigma Aldrich) was added. The gel was let solidify for 20 min, after which the isopropanol layer was removed. A solution of 155 mM Tris-HCl (pH=6.5), 10 v/v% 40% acrylamide solution (Biorad), 0.2 w/v% sodium dodecyl sulfate (Sigma Aldrich), 0.1 w/v% ammonium persulfate (Sigma Aldrich), and 0.1 v/v% N,N,N’,N’-tetramethyl ethylenediamine (Sigma Aldrich) was prepared and added to the existing gel layer, after which a well comb (Biorad) was added. The gel was let solidify for 20 min.

Protein samples were mixed with SDS loading buffer (Laemmli buffer, [Laemmli, 1970]) and kept for 5 min at 95^°^C before loading onto the gels. Precision Plus Protein Unstained standard (Biorad) (and Precision Plus Protein All Blue Standard (Bio-rad) in case of Western blot analysis) was used as a protein standard. Gels were run for 5 min at 130 V followed by 55min min at 180 V (PowerPac Basic Power Supply (Biorad)).

If not stated otherwise, imaging was done on a ChemiDoc MP (Biorad) using the ‘Stain-free gels’ feature and automatic exposure time determination. In some instances, gels were stained using SimplyBlue SafeStain (Invitrogen), SYPRO Ruby protein gel stain (Invitrogen), or SilverQuest Staining Kit (Invitrogen) following the basic steps of the accompanying manual. Imaging was done on a ChemiDoc MP (Biorad) using automatic exposure time determination.

### Western blotting

After SDS-PAGE, the sample was transferred from the SDS-PAGE to a blotting membrane (Trans-Blot Turbo Transfer Pack, Bio-rad) using the ‘Mixed MW’ program of the Trans-Blot Turbo Transfer System (Bio-rad). The blotting membrane was incubated with Immobilon signal enhancer (Millipore) at room temperature for 1 h. The blotting membrane was incubated with primary antibody (His Tag Antibody, Mouse (OAEA00010, Aviva Systems Biology) (dilution: 1:4000)), diluted in Immobilon signal enhancer, at room temperature for 1 h. It was washed thrice with TBS-T (10 mM Tris-HCl (pH=7.5), 150 mM NaCl, 0.1 v/v% Tween-20 (Sigma Aldrich)). For each washing step the blotting membrane was incubated with TBS-T at room temperature for 20 min. The blotting membrane was incubated with secondary antibody (IgG2b Antibody HRP conjugated (Goat Anti-Mouse) (OASA06620, Aviva Systems Biology) (dilution: 1:1000)), diluted in Immobilon signal enhancer, at room temperature for 1 h, after which it was again washed thrice with TBS-T. SuperSignal West Pico Mouse IgG Detection Kit (Thermo Scientific) was used for activation. Imaging was done on a ChemiDoc MP (Biorad) using the ‘Chemi Sensitive’ feature and automatic exposure time determination.

## Abbreviations

aa: amino acid
BSA: Bovine serum albumin
GAP: GTPase activating protein
GEF: GDP/GTP exchange factor
SEC-MALS: size exclusion chromatography - multi-angle light scattering

## Data availability

The data underlying this publication are openly available at https://data.4tu.nl at doi.org/10.4121/56535d83-8a7a-4367-9ae5-f0d56b51161f (CC BY-SA 4.0) [Tschirpke et al., 2025].

## Acknowledgements

We thank R. van der Valk and C. de Agrela Pinto for experimental assistance, F. Shamsi and S. Farooq for their pioneering work of GTPase activity by Cdc42 and its regulators, and M. Depken for discussions and careful reading of the manuscript. We thank D. McCusker (University of Bordeaux) for the plasmid pDM272 and N. Dekker (TU Delft) for the plasmid pET28a-His-mcm10-Sortase-Flag.

## Contributions

**S. Tschirpke:** Conceptualization, Methodology, Investigation, Formal analysis, Validation, Writing - Original Draft, Writing - Review & Editing, Visualization, Project administration. **W. K.-G. Daalman:** Conceptualization, Methodology, Investigation, Software, Formal analysis, Validation. **F. van Opstal:** Investigation, Formal analysis, Validation. **L. Laan:** Conceptualization, Methodology, Investigation, Writing - Review & Editing, Funding acquisition, Project administration.

## Supplement S1

We developed a Cdc42 GTPase activity model for determining the GTPase cycling rates *k*. The model is described in detail in [Tschirpke et al., 2024]. Key aspects are summarized here:

### GTPase model

Cdc42 GTPase cycling involves three steps: (1) A GTP molecule from solution binds to Cdc42. (2) Cdc42 hydrolyses GTP. (3) Cdc42 releases GDP.

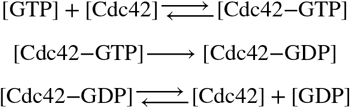

It can further be upregulated by effector proteins: GAPs have been shown to enhance GTP hydrolysis by Cdc42 (step 2), GEFs enhance the release of GDP from Cdc42 (step 3) [Park and Bi, 2007, Martin, 2015, Chiou et al., 2017].

To quantitatively describe the GTPase reaction cycle, we coarse-grained the GTPase reaction steps with

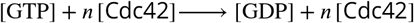

To account for possible Cdc42 dimerization and cooperativity, we included the following reactions into the model:

1. We assume that Cdc42 can dimerize, as other small GTPases have been shown to dimerize [Zhang and Zheng, 1998, Zhang et al., 1999, Zhang et al., 2001, Kang et al., 2010]:

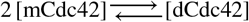

and both monomeric and dimeric Cdc42 can contribute to the overall GTP hydrolysis with different rates:

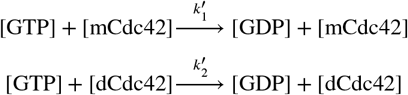 Assuming that the majority of Cdc42 is in its monomeric form ([mCdc42] < *C*_*d*_, with *C*_*d*_ as the concentration at which half of the total Cdc42 is dimeric), we can approximate

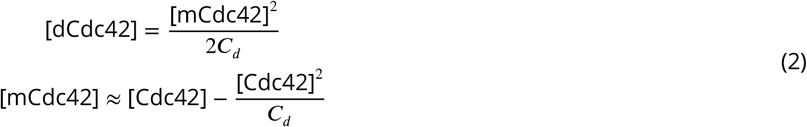
2. Next to cooperativity from dimerization, cooperativity can also emerge when Cdc42 proteins come in close contact with each other - they can affect each other’s behavior without forming a stable homodimer, effectively functioning as an effector protein for themselves:

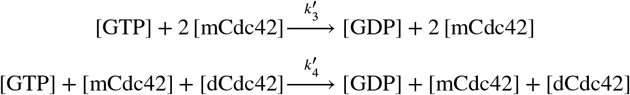
3. Effector proteins, such as GAPs and GEFs, affect the speed of the GTP hydrolysis cycle:

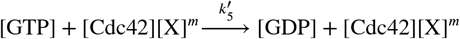

Here X is an effector protein.

Our data shows that the amount of remaining GTP follows an exponential decline over time (S1 Fig. 1):

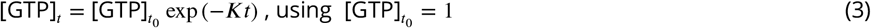

Considering reactions (1) - (3), we can thus define *K* in Eq. 3 as

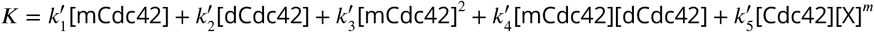

Using Eq. 2, and considering only up to second-order terms, results in

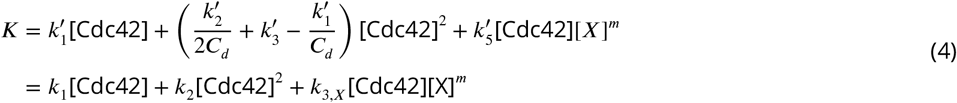

where *k*_1_ refers to GTP hydrolysis cycling rates of monomeric Cdc42, *k*_2_ includes effects of cooperativity and dimerization and *k*_3_ represents the rate of Cdc42 - effector interaction. We refer to *K* as ‘overall GTP hydrolysis rate’.

### Variability between assays

We used Eq. 4 with [X]=0 to determine the rates of Cdc42 alone. We then conducted assays with Cdc42 and an effector protein to determine *k*_3_. While doing so we needed to account for assay variability, i.e. for the observation that the rates for Cdc42 can vary between assays. Possible reasons for this include small concentration differences introduced though pipetting of small volumes (as are required for this assay), temperature and shaker speed fluctuations during the incubation step, and/or intrinsic changes in the protein activities due to other external conditions. To account for this variance, we introduced the parameter *c*_*corr*_. It maps all factors that lead to variations between assays onto the Cdc42 concentration.

The assay data, including samples containing only Cdc42 and Cdc42 - (effector) protein mixtures, was fitted with

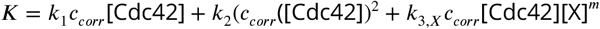

to determine *c*_*corr*_ and *k*_3,*X*_ (using *k*_1_ and *k*_2_ determined earlier).

Only assays with *c*_*corr*_ values from 0.5 to 1.5 were used for analysis. Generally, with *c*_*corr*_ values were distributed around 1.0, confirming that the variation between assays is small (S1 Fig. 2).

### Pooled estimates

The pooled estimates of rates *k*_1_, *k*_2_, and *k*_3_ were determined through weighting their standard error, as described in the following:

Within an assay, the rate parameters per run are calculated, but also a weighted average can be taken from these values to create a pooled estimate. Concretely:

For pooling, we model the *n* parameter estimates *y*_*i*_ to originate from a single pooled estimate *a* as:

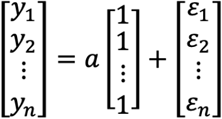

with *ϵ*_*i*_ *∼ N*(0, *σ*_*i*_) where *σ*_*i*_ is the standard error of the parameter estimate *y*_*i*_. As uncertain estimates should be weighted less, the natural weights *w*_*i*_ to each *y*_*i*_ should be 1/*σ*_*i*_, after which all weighted errors follow a standard normal distribution:

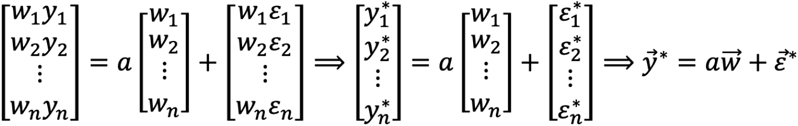

Getting the estimate for *a*, namely *â*, is the result of a simple regression (i.e. weighted least squares), minimizing the sum of squared errors (see e.g. [Heij et al., 2004]):

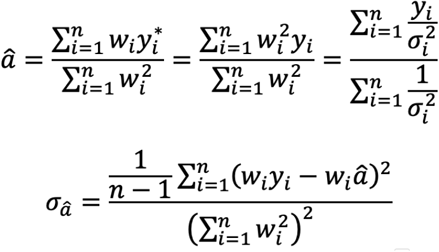

**S1 Figure 1.**
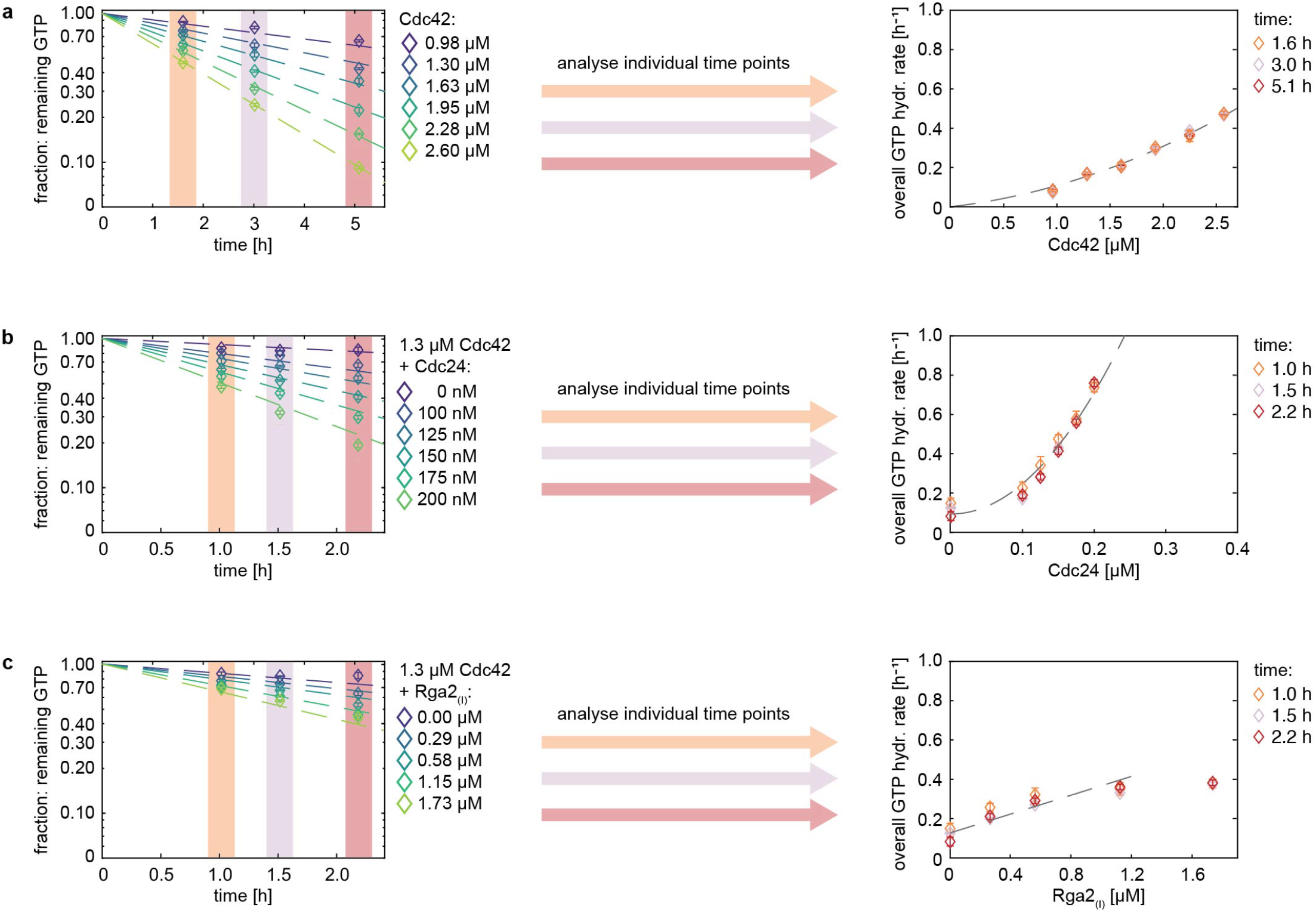
The GTP concentration declines exponentially with time in GTPase reactions. Amount of remaining GTP for (a) Cdc42 concentrations (b) Cdc42 - Cdc24 mixtures, and (c) Cdc42 - Rga2_(*I*)_ mixtures, each for three time points (measured as one individual assay per time point). The remaining GTP content declines exponentially with time (left). Data of each individual time point shows the same overall GTP hydrolysis cycling rate for each GTPase - effector mixture. Thus, only one time point per assay condition is needed, to fit the data (right).

**S1 Figure 2.**
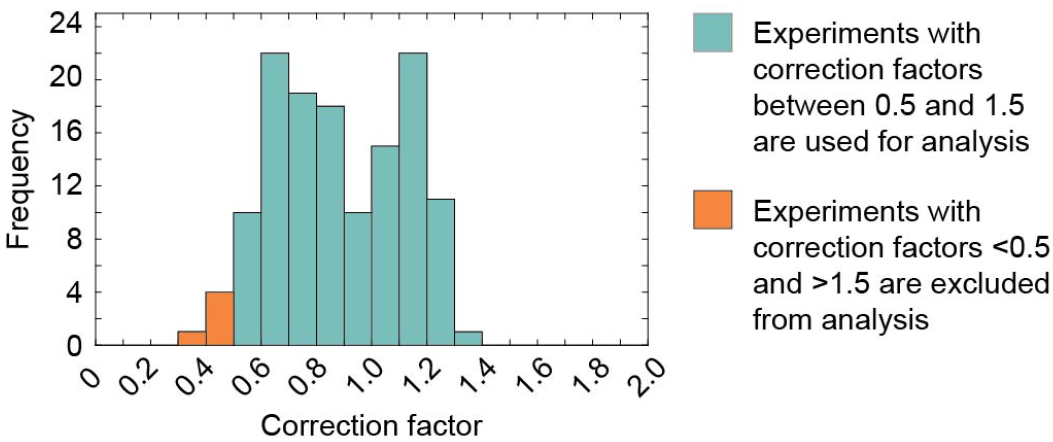
The variation between GTPase assays is small. *c*_*corr*_ values are distributed around 1.0.

## Supplement S2

We found that Cdc24’s effect of the overall GTP hydrolysis rate *K* increases non-linearly with its concentration (Fig. 3b), which we speculate to be linked to Cdc24 di-or oligomerization: Cdc24 has the capability to oligomerize via its DH domain [Mionnet et al., 2008]. Dimers and oligomers could have an increased GEF activity through releasing Cdc24 from its self-inhibited state [Shimada et al., 2004].

This hypothesis is in contrast with *in vitro* work on peptides based on Cdc24 fragments, showing that these peptides exhibit an oligomerization-independent GEF activity [Mionnet et al., 2008]. These findings would exclude that Cdc24 oligomers exhibit an increased GEF activity.

However, Mionnet *et al*. used peptides that were based on some of Cdc24’s domains (DH and PH domain, aa 285-681), and not full-length protein, for their *in vitro* GEF activity assays. Other domains that are not directly involved in oligomerization or GEF function can still affect these properties. For example, the PB1 domain was suggested to reduce Cdc24 GEF activity in a self-inhibitory fashion [Shimada et al., 2004]. Next, oligomerization was induced through a chemical; the peptides contained an additional oligomerization domain (FKBP) that was not related to Cdc24 and could be triggered to oligomerize through the addition of a chemical (synthetic oligomerization inducer AP20187). Thus, the findings on Cdc24 peptides might not fully translate to full-length Cdc24.

To investigate our hypothesis of an oligomerization-induced increase of Cdc24’s GEF activity, we investigated two Cdc24 mutants: **Cdc24-DH3** (mutations L339A and E340A) with a reported 2.5× reduction in oligomerization and **Cdc24-DH5** (mutation F322A) with a reported 10× reduction in oligomerization [Mionnet et al., 2008].

For clarity, we like to note that the oligomerization studies conducted by Mionnet *et al*. utilized full-length Cdc24, while the *in vitro* GEF activity assays utilized peptides based on Cdc24 fragments and chemically induced oligomerization. While we question the generality of an oligomerization-independent GEF activity of Cdc24 (obtained using Cdc24 peptides and chemically induced oligomerization), we do not question their findings on Cdc24 oligomerization (obtained using full-length protein).

We recombinantly expressed Cdc24-DH3 and Cdc24-DH5 and purified them using His-affinity chromatography. Cdc24-DH3 expressed as the full-length protein, while for Cdc24-DH5 only fragments remained (S2 Fig. 1). Mionnet *et al*. had already observed that *S. cerevisiae* cells expressing Cdc24-DH5 as sole copy were not viable and suggested that this was due to Cdc24-DH5’s inability to localize correctly [Mionnet et al., 2008]. Although our data does not disprove this conclusion, it presents an additional possibility: If the DH5 mutation disrupts Cdc24 folding and expression, cells expressing Cdc24-DH5 might not be viable because they lack full-length Cdc24.

We continued our investigation utilizing Cdc24-DH3 (which was further purified using size exclusion chromatography, S11): We conducted GTPase assays to determine the concentration-dependent increase of the overall GTP hydrolysis rate *K* through Cdc24-DH3 (Fig. 3c), fitting the data with our standard exponential model (Fig. 3d). Through comparing rates *k*_3_, we found that Cdc24-DH3 has a 17× reduced GEF activity compared to wildtype Cdc24 (Fig. 3e, Tab. 2). Cdc24-DH3 contains mutations L339A and E340A, which are located in the Dbl homology (DH) domain. The DH domain is linked to GTPase interaction and activation, and responsible for Cdc24 oligomerization. It is thus possible that (1) the mutations lead to both a reduction in oligomerization and a reduced GEF activity, or (2) the mutations lead to reduced oligomerization, resulting in a reduced GEF activity. However, given that peptides comprising Cdc24’s DH domain did not show a reduced GEF activity when the DH3 mutations were introduced [Mionnet et al., 2008], we believe that the reduced GEF activity of full length Cdc24-DH3 stems from its reduced ability to oligomerize.

We hypothesized that the non-linear rate increase through Cdc24 stems from Cdc24 oligomerization: Cdc24 oligomers could have an increased GEF activity through being released of the self-inhibited state Cdc24 monomers are in [Shimada et al., 2004]. With increasing Cdc24 concentration the amount of Cdc24 oligomers increases, resulting in a non-linear increase of the overall GTP hydrolysis rate *K*. Cdc24-DH3 has a 2.5× reduction in its oligomerization capacity [Mionnet et al., 2008]. If the non-linear rate increase through Cdc24 is indeed linked to its oligomerization, Cdc24-DH3 should show a more linear (/less quadratic) rate increase. We therefore fitted data of both Cdc42 - Cdc24 and Cdc42 - Cdc24-DH3 mixtures, only using assays with at least five data points, with a modified version of our model:

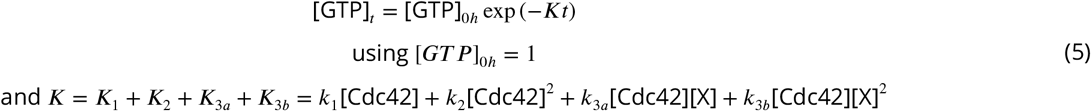

Here *X* is either Cdc24 or Cdc24-DH3. *K*_3*a*_ scales linearly with the Cdc24/Cdc24-DH3 concentration, and *K*_3*b*_ scales quadratically with the Cdc24/Cdc24-DH3 concentration. We determined the ratio of the two rates (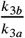, S2 Tab. 1). The larger this ratio, the more quadratic (less linear) the overall GTP hydrolysis rate *K* increases with Cdc24/Cdc24-DH3 concentration. Cdc24-DH3 data exhibits 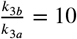 and Cdc24 data exhibits 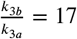, suggesting that the rate increase through Cdc24-DH3 is indeed more linear than the rate increase through wildtype Cdc24. These findings support our hypothesis that the non-linear rate increase of the overall GTP hydrolysis rate *K* through Cdc24 might originate from Cdc24 oligomerization.

Taken together, our data suggests that

1. **Cdc24’s GEF activity might be oligomerization-dependent**. Our data on the Cdc24-DH3 mutant, which has a 2.5× reduced oligomerization capacity [Mionnet et al., 2008], shows that: (1) Cdc24-DH3 has a reduced GEF activity. (2) The overall GTP hydrolysis rate *K* increases more linearly (/less quadratically) with Cdc24-DH3 concentration than with the concentration of wildtype Cdc24. Both findings support our hypothesis that Cdc24’s GEF activity is oligomerization-dependent.
2. **Findings obtained using peptides of Cdc24 fragments do not translate to full-length Cdc24**. Mionnet *et al*. found that a chemically induced oligomerization of peptides based on Cdc24 fragments does not affect their GEF activity [Mionnet et al., 2008]. We found that a Cdc24 mutant with reduced oligomerization capacity has a reduced GEF activity, questioning the generalizability of data based on protein fragments.

**S2 Figure 1.**
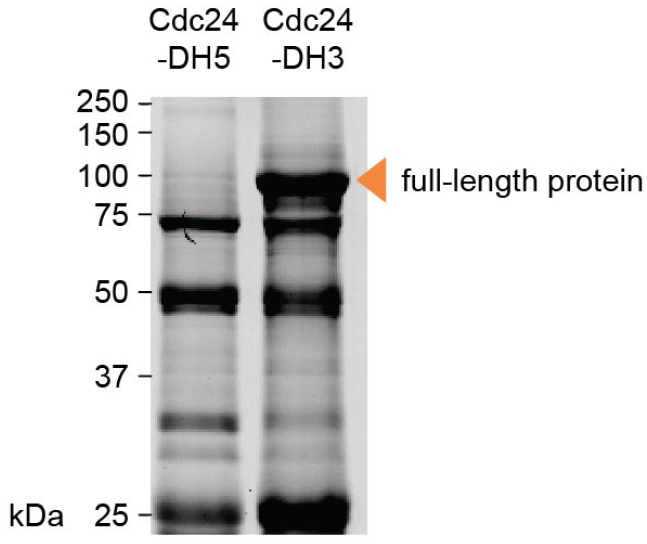
Cdc24-DH3 expression yielded the full-length protein, whereas for Cdc24-DH5 only fragments could be detected. SDS-PAGE of Cdc24 mutants Cdc24-DH5 and Cdc24-DH3 after His-affinity chromatography.

**S2 Table 1.** Interaction rates *k*_3*a*_ and *k*_3*b*_ of Cdc42 - Cdc24/Cdc24-DH3 mixtures (determined using Eq. 5).

| | $k_{3a}$<br>[$\mu\text{M}^{-3} \text{ h}^{-1}$] | $k_{3a}$ std. err.<br>[$\mu\text{M}^{-3} \text{ h}^{-1}$] | $k_{3b}$<br>[$\mu\text{M}^{-4} \text{ h}^{-1}$] | $k_{3b}$ std. err.<br>[$\mu\text{M}^{-4} \text{ h}^{-1}$] | $k_{3b}/k_{3a}$<br>[$\mu\text{M}^{-1} \text{ h}^{-1}$] |
| --- | --- | --- | --- | --- | --- |
| Cdc24: pooled estimate (n=16) | 0.409 | 0.049 | 6.768 | 0.191 | 17 |
| Cdc24-DH3: pooled estimate (n=4) | 0.069 | 0.014 | 0.663 | 0.083 | 10 |

## Supplement S3

**S3 Figure 1.**
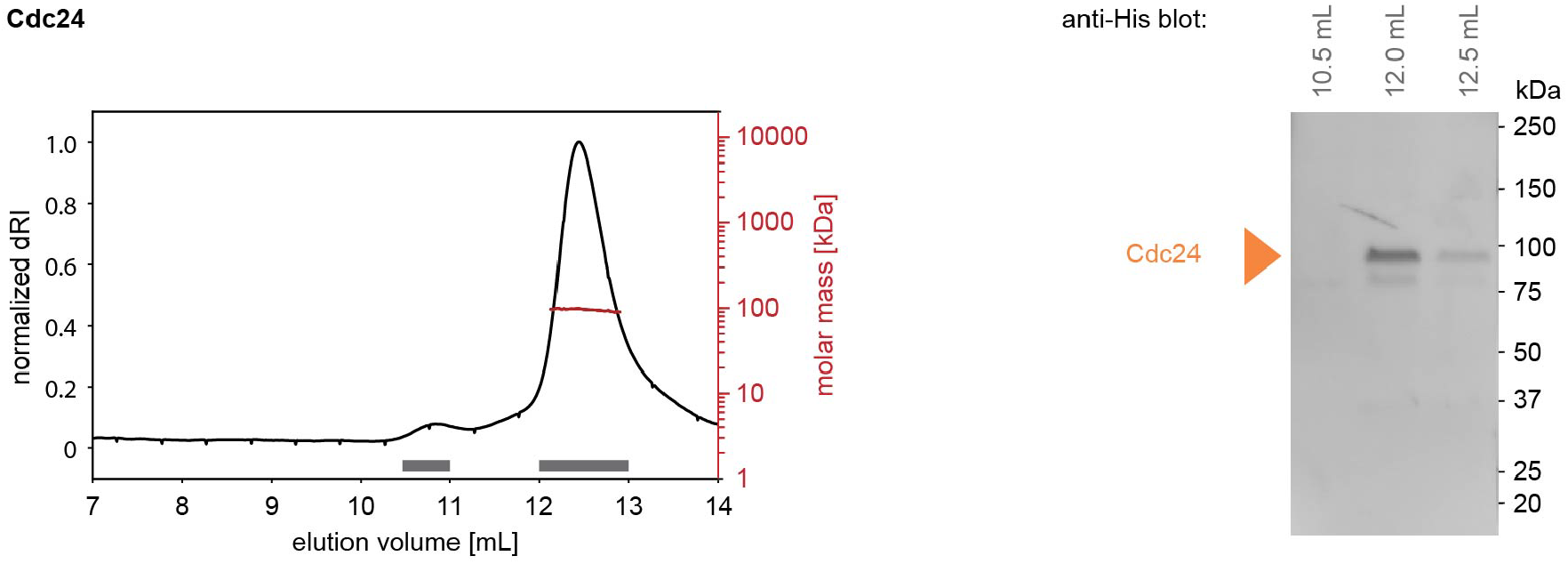
SEC-MALS analysis of Cdc24. Size-exclusion profile and MALS analysis (left) and Western blot analysis of SEC-MALS elution fractions (right).

## Supplement S4

To ensure the effects we observed in our assays are protein-specific, we conducted assays with bovine serum albumin (BSA) and Casein - two proteins considered inert. These proteins are expected to not affect the assay or interact with GTPases. In absence of GTPases both BSA and Casein did not lead to hydrolyzed GTP, showing that they do not cause GTP hydrolysis themselves and that they do not affect any downstream processes of the assay (S4 Fig. 1). They slightly increase the overall GTP hydrolysis rate of Cdc42 (S4 Fig. 2) and to a smaller extend that of Ras (S4 Fig. 3). The BSA and Casein concentrations used here (0-5 µM) are far below concentrations where crowding effects are expected [Chebotareva et al., 2004], excluding crowding effects. We suspect that through sticking to reaction chamber walls BSA and Casein increase the effective GTPase concentration in the assay, thus causing a slight increase in the overall GTP hydrolysis rate. Supporting this line of reasoning is the observation that the effect of Casein on Cdc42 is larger than its effect on Ras: Cdc42 has a higher GTPase activity than Ras (S4 Tab. 1). An effective increase in Cdc42’s effective concentration will thus have a larger effect on it’s overall GTP hydrolysis rate *K* than a similar concentration increase of Ras. The effect of BSA and Casein is at least 3.7× smaller than the effect of Rga2_(*I*)_ (the weakest of Cdc42’s effectors) (S4 Tab. 2), suggesting that their non-canonical effect on the assay does not play a major role.

We conducted some preliminary assays with (a) Cdc42 - Cdc24 - BSA and (b) Cdc42 - Rga2_(*I*)_ - BSA mixtures (S4 Fig. 4): In our model the effectors contribute to the overall GTP hydrolysis rate *K* of Cdc42 through three terms: *K*_3,*X*1_, *K*_3,*X*2_, and *K*_3,*X*1,*X*2_ where *X*1 and *X*2 are (a) Cdc24 and BSA and (b) Rga2_(*I*)_ and BSA. *K*_3,*Cdc*24_, *K*_3,*Rga*2_, and *K*_3,*BSA*_ represent the rate contribution of Cdc24, Rga2_(*I*)_, and BSA alone. They are in the three-protein mixture (Cdc42 - Cdc24 - BSA, Cdc42 - Rga2_(*I*)_ - BSA) roughly the same as when Cdc42 was incubated with one effector alone (Cdc42 + Cdc24, Cdc42 + Rga2, or Cdc42 + BSA) (S4 Fig. 4c). The interaction term *K*_3,*Cdc*24,*BSA*_ of Cdc24 and BSA was roughly the same as the individual contribution of Cdc24 (*K*_3,*Cdc*24,*BSA*_ ≈ *K*_3,*Cdc*24_) and Rga2 and BSA showed no interaction (*K*_3,*Rga*2,*BSA*_ ≈ 0)^2^ (S4 Tab. 3). If BSA is sticking to reaction chamber walls, it can increase the effective concentration of Cdc42 and Cdc24/Rga2 in the assay. Given that Cdc24 had a strong and Rga2 had a weak effect on Cdc42, an increase in the effective Cdc24 concentration results in a significant increase of the overall GTP hydrolysis rate (leading to a positive *K*_3,*Cdc*24,*BSA*_), whereas an increase in the effective Rga2 concentration has almost no observable effect (*K*_3,*Rga*2,*BSA*_ ≈ 0). Importantly, *K*_3,*Cdc*24,*BSA*_ is seven times smaller than *K*_3,*Cdc*24,*Rga*2_ (the interaction term between Cdc24 and Rga2_(*I*)_ in Cdc42 - Cdc24 - Rga2_(*I*)_ assays, Fig. 5), confirming that the observed synergy between Cdc24 and Rga2_(*I*)_ is not due to non-canonical effects.

**S4 Figure 1.**
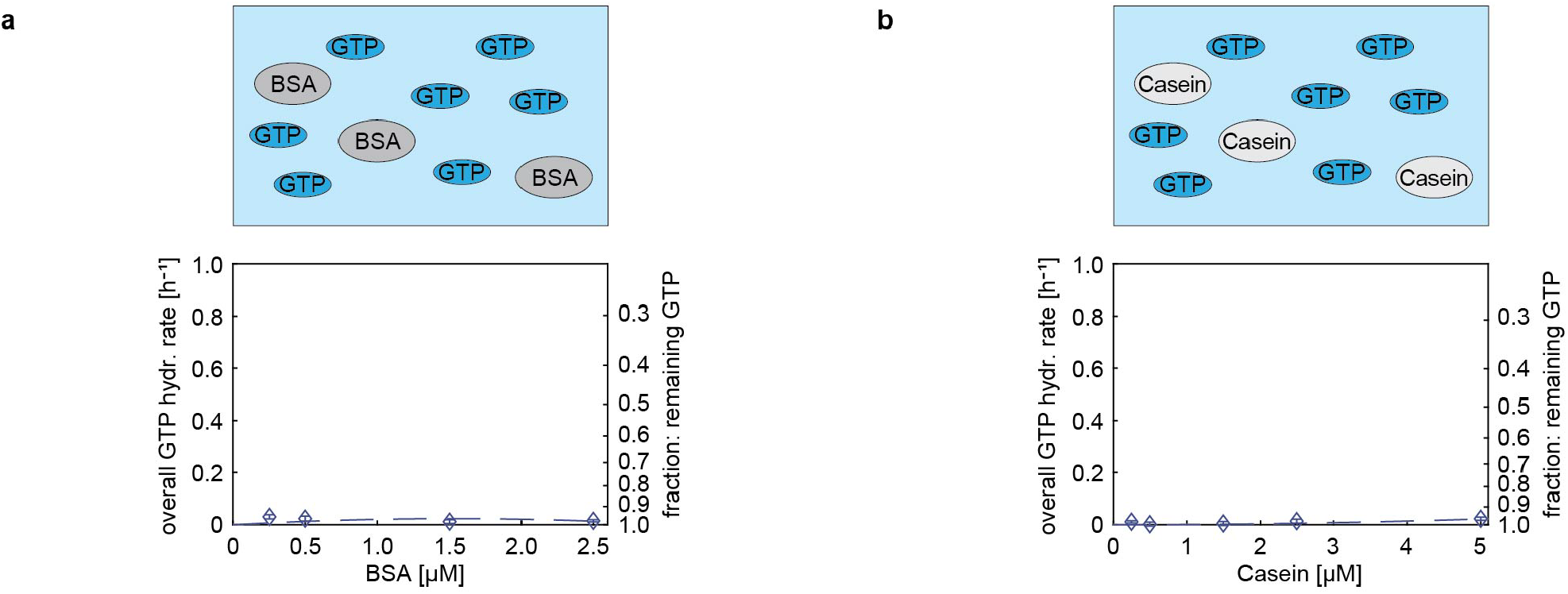
In absence of an GTPase enzyme, BSA and Casein do not lead to GTP hydrolysis (or affect any downstream processing steps of the assay). (a) BSA incubated with GTP. (b) Casein incubated with GTP.

**S4 Figure 2.**
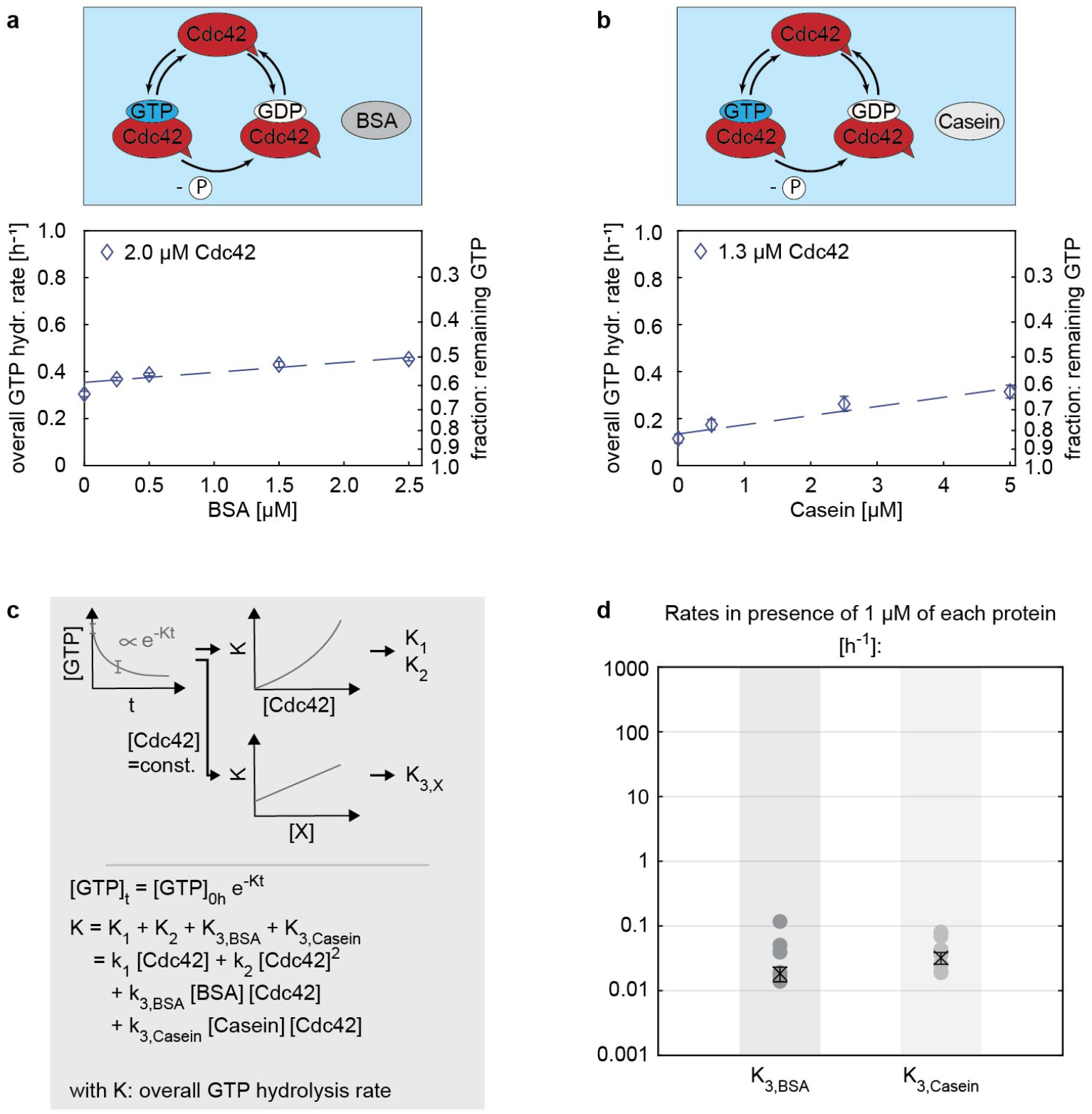
Non-canonical effects in GTPase assays: inert proteins (BSA, Casein) slightly boost Cdc42’s GTPase activity. (a,b) The overall GTP hydrolysis rate *K* of Cdc42 in presence of varying BSA (a) and Casein (b) concentrations. The increase in Cdc42’s overall GTP hydrolysis rate could be due to (1) unknown non-specific interactions or (2) because BSA and Casein coat to the reaction chamber walls, preventing Cdc42 from binding, thus increasing the active Cdc42 concentration in the reaction chamber. (c) Illustration of the data processing and fitting model. (d) Summary of the rates *K*. The values shown refer to the rate values in presence of 1 µM of each protein, e.g. ‘*K*_3,*BSA*_’ refers to ‘*k*_3,*BSA*_[BSA][Cdc42]’ with [Cdc42]=[BSA]=1 µM. Crosses with error bars represent the weighted mean and the standard error of the mean (S1), filled circles show individual measurements.

**S4 Figure 3.**
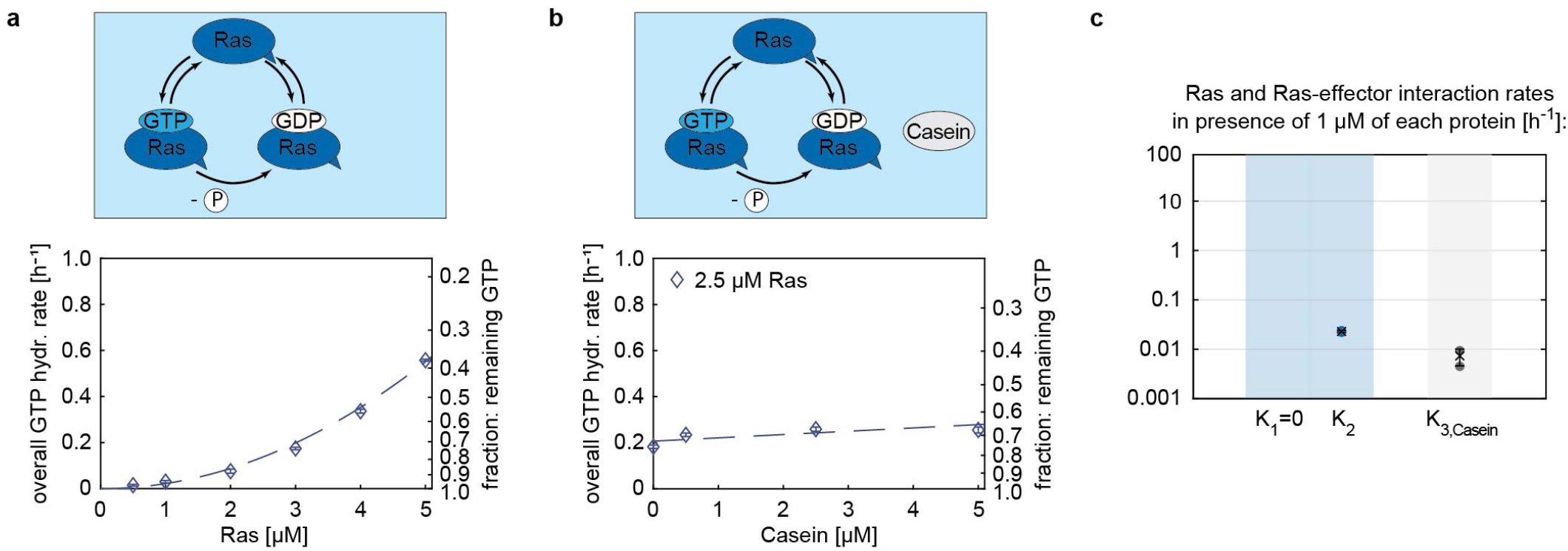
Non-canonical effects in GTPase assays: the inert protein Casein slightly boosts the GTPase activity of Ras. (a) The overall GTP hydrolysis rate (*K*) of Ras. (b) The overall GTP hydrolysis rate (*K*) of Ras in presence of varying Casein concentrations. The increase in the overall GTP hydrolysis rate could be due to (1) unknown non-specific interactions or (2) because Casein coats to the reaction chamber walls, preventing Ras from binding, thus increasing the active Ras concentration in the reaction chamber. (c) Summary of the rates *K*. The values shown refer to the rate values in presence of 1 µM of each protein, e.g. ‘*K*_3,*Casein*_’ refers to ‘*k*_3,*Casein*_[Casein][Ras]’ with [Ras]=[Casein]=1 µM. Crosses with error bars represent the weighted mean and the standard error of the mean (S1), filled circles show individual measurements.

**S4 Figure 4.**
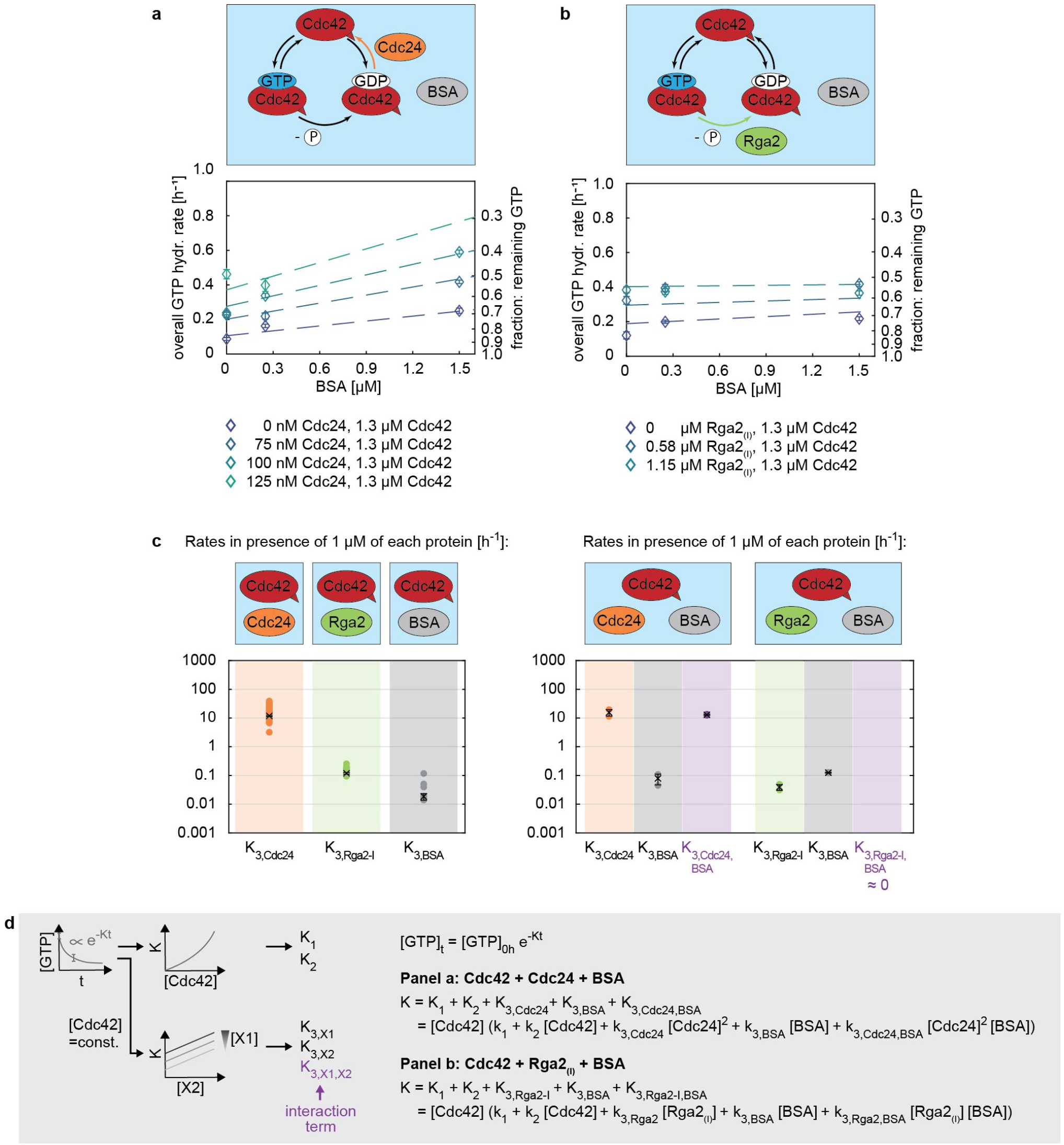
Non-canonical effects in GTPase assays: the inert proteins BSA slightly enhances the effect of Cdc24, but not of Rga2_(*I*)_, on Cdc42’s GTPase activity. (a,b) Increase in the overall GTP hydrolysis cycling rate *K* of Cdc42 in presence of (a) Cdc24 and BSA, and (b) Rga2 and BSA. (c) Summary of the rates *K* obtained in the three-protein assay (right) in comparison to those of the two-protein assay (left): In three-protein assays the rate contribution of the individual proteins is comparable to those obtained in the two-protein assay. Additionally, an interaction rate is obtained (shown in purple). In the case of Cdc42-Cdc24-BSA and Cdc42-Rga2-BSA mixtures, this interaction rate is comparable to the Cdc24 contribution/zero, indicating a weak/ almost no synergy. This effect could be due to non-specific protein-protein binding or because BSA coats to the reaction chamber walls, preventing the other proteins from binding, thus increasing their active concentration in the reaction chamber. The values shown refer to the rate values in presence of 1 µM of each protein, e.g. ‘*K*_3,*BSA*_’ refers to ‘*k*_3,*BSA*_[BSA][Cdc42]’ with [Cdc42]=[BSA]=1 µM. Crosses with error bars represent the weighted mean and the standard error of the mean (S1), filled circles show individual measurements. (d) Illustration of the data processing and fitting model.

**S4 Table 1.** GTP hydrolysis cycling rates *k*_1_ and *k*_2_ of Cdc42 and Ras.

| | $k_1$<br>[$\mu\text{M}^{-1} \text{h}^{-1}$] | $k_1$ std. err.<br>[$\mu\text{M}^{-1} \text{h}^{-1}$] | $k_2$<br>[$\mu\text{M}^{-2} \text{h}^{-1}$] | $k_2$ std. err.<br>[$\mu\text{M}^{-2} \text{h}^{-1}$] |
| --- | --- | --- | --- | --- |
| Ras, pooled estimate (n=2) | 0.000 | 0.000 | 0.021 | 0.001 |
| Cdc42, pooled estimate (n=5) | 0.110 | 0.002 | 0.013 | 0.001 |

**S4 Table 2.**
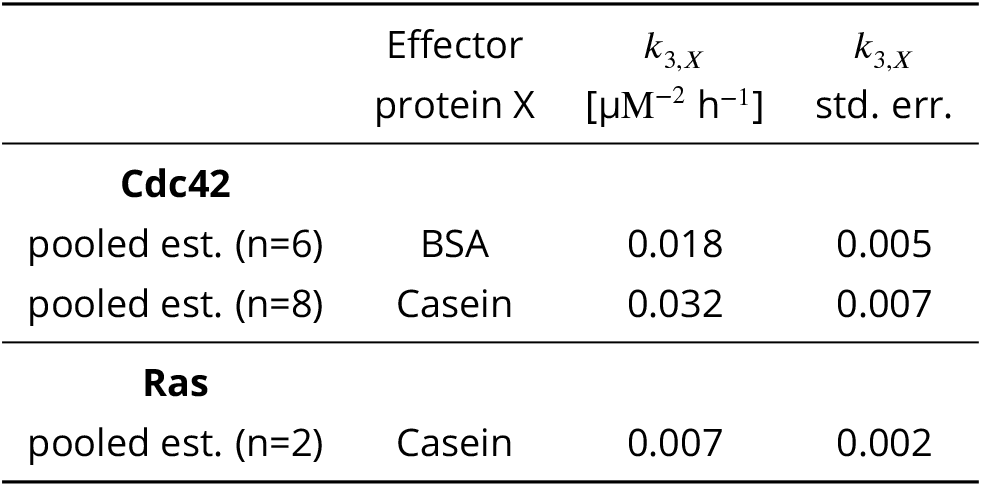
Interaction rates *k*_3,*X*_ of GTPase - effector protein mixtures.

**S4 Table 3.** Cdc42 - effector protein *X*_1_ - effector protein *X*_2_ interaction rates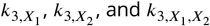. *: unit in case of X_1_=Cdc24: [µM^−3^ h^−1^]. **: unit in case of X_1_=Cdc24: [µM^−4^ h^−1^].

| | Effector<br>protein $X_1$ | Effector<br>protein $X_2$ | $k_{3,X_1}$<br>[$\mu\text{M}^{-2} \text{h}^{-1}$] * | $k_{3,X_1}$<br>std. err. | $k_{3,X_2}$<br>[$\mu\text{M}^{-2} \text{h}^{-1}$] | $k_{3,X_2}$<br>std. err. | $k_{3,X_1,X_2}$<br>[$\mu\text{M}^{-3} \text{h}^{-1}$] ** | $k_{3,X_1,X_2}$<br>std. err. |
| --- | --- | --- | --- | --- | --- | --- | --- | --- |
| pooled est. (n=7) | Cdc24 | Rga2<sub>(I)</sub> | 15.126 | 0.736 | 0.131 | 0.015 | 86.961 | 6.055 |
| pooled est. (n=2) | Cdc24 | BSA | 15.642 | 4.054 | 0.079 | 0.484 | 12.887 | 0.538 |
| pooled est. (n=2) | Rga2<sub>(I)</sub> | BSA | 0.125 | 0.166 | 0.039 | 0.009 | -0.024 | 0.003 |

## Supplement S5

Fitting GTPase assays data where both the GEF Cdc24 and the GAP Rga2 where added resulted in a positive synergy term (Fig. 5), which could originate from two sources: (1) Cdc24-Rga2 synergy due to rate-limiting effects of the GTPase cycle, and (2) Cdc24-Rga2 synergy due to protein-protein interactions.

In the following we describe the rate-limiting step model and then discuss to which extend the rate-limiting step model explains our data and thus the positive synergy term.

### The rate-limiting step model

In the rate-limiting step model, we assume that we have a GTPase cycle in which at least one of the three GTPase cycle steps is rate-limiting: (A) GTP binding, (B) GTP hydrolysis, and (C) GDP release. We assume that the addition of a GEF accelerates only GDP release (step C), and that the addition of a GAP only accelerates GTP hydrolysis (step B).

Biochemically, all three steps in the GTPase cycle are expected to be relevant. However, here we will consider only the final two steps, as sensitivity to rate limitation by GAP/GEF is maximized when time spent in the GAP/GEF-independent step in the cycle (step A: GTP binding) is negligible (i.e. never rate-limiting).

The rate-limiting step model thus consists of two steps:

1. a nucleotide exchange step (step C+A) which is dominated by GDP release (step C) and assumed to be accelerated exclusively by the GEF, and
2. a GTP hydrolysis step (step B) exclusively enhanced by the GAP.

In the rate-limiting step model, addition of the GEF to the GTPase would accelerate only step (1), leaving a slow GTP hydrolysis step (step 2) that limits the overall cycling rate. Analogously, addition of GAP would accelerate only step (2), leaving a nucleotide exchange step (step 1) that limits the overall cycling speed. Addition of both GEF and GAP would speed up both steps, and therefore synergistically accelerate the overall cycling rate. The synergy is then a property of the GTPase cycle, and not due to proteins enhancing each other’s activity.

Specifically, GEF-GAP synergy can appear if one of the two conditions applies:

1. the addition of a GEF speeds up the GDP release step (step 1) so much that the release step stops (or almost stops) being the rate-limiting step, or
2. the addition of a GAP speeds up the GTP hydrolysis step (step 2) so much that the hydrolysis step stops (or almost stops) being the rate-limiting step.

In these conditions, the acceleration of the GTPase cycle, accomplished by adding only a GEF or adding only a GAP, is interdependent. Therefore, we **consider the possible acceleration of the GTPase cycle by GAP and GEF individually** (i.e. assays comprising of Cdc42 + GEF and assays comprising of Cdc42 + GAP), and compare these to our observations to determine whether the rate-limiting step model can fully explain our data.

The GTPase cycle time *T*_*c*_ (with the rate *r*_*c*_) is thus composed of hydrolysis time *T*_*h*_ and nucleotide exchange time *T*_*e*_, and the respective rates *r*_*h*_ and *r*_*e*_, are connected through:

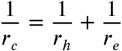

If we compare the ratio of the rates with one effector (GEF or GAP) added in the assay (index 1) with the basal rate without an effector added (i.e. Cdc42 only) (index 0), we obtain the cycle acceleration factor *a*:

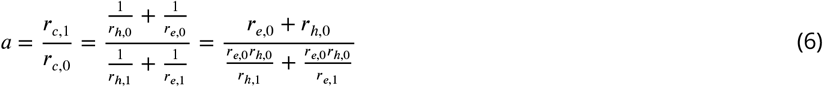

Using

- *r*_*c*,0_: the basal GTPase cycle rate (of only Cdc42)
- *r*_*c*,1_: the GTPase cycle rate with an effector added (GEF or GAP)
- *r*_*h*,0_: the basal GTP hydrolysis rate (of only Cdc42)
- *r*_*h*,1_: the GTP hydrolysis rate with an effector added (GEF or GAP)
- *r*_*e*,0_: the basal nucleotide exchange rate (of only Cdc42)
- *r*_*e*,1_: the nucleotide exchange rate with an effector added (GEF or GAP)

There is an interdependence between how much the GAP and the GEF can accelerate the GTPase cycle, if the GAP and GEF are assumed to only accelerate GTP hydrolysis and nucleotide exchange respectively. E.g., how much the total GTPase cycle rate *r*_*c*_ is accelerated by an increase in the GTP hydrolysis rate *r*_*h*_ depends on and can be limited by the current nucleotide exchange rate *r*_*e*_.

When we **only add a GEF** and the GEF accelerates only the nucleotide exchange rate *r*_*e*,1_ (and not the GTP hydrolysis rate *r*_*h*,1_, meaning *r*_*h*,1_ = *r*_*h*,0_), then the maximal total GTPase cycle rate acceleration *a*_*GEF*_ that the GEF can accomplish is when

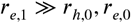

I.e. when the nucleotide exchange rate in presence of a GEF *r*_*h*,1_ is much bigger than the basal hydrolysis rate *r*_*h*,0_ and much bigger than the basal nucleotide exchange rate *r*_*e*,0_:

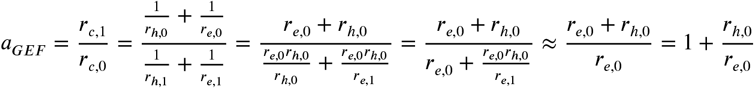

We thus assume the GEF accelerates the cycle so much that the nucleotide exchange step is much faster than the hydrolysis step, at which point the effect of adding more GEF would saturate.

Analogously, when we **only add a GAP** and the GAP accelerates only the GTP hydrolysis rate *r*_*h*,1_ (and not the nucleotide exchange rate *r*_*e*,1_, meaning *r*_*e*,1_ = *r*_*e*,0_), then the maximal total GTPase cycle rate acceleration *a*_*GAP*_ that the GAP can accomplish is when

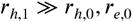

i.e. when the hydrolysis rate in presence of a GAP *r*_*h*,1_ is much bigger than the basal hydrolysis rate *r*_*h*,0_ and much bigger than the basal nucleotide exchange rate *r*_*e*,0_:

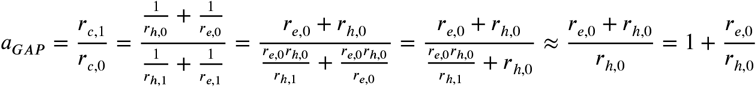

We thus assume the GAP accelerates the cycle so much that the hydrolysis step is much faster than the nucleotide exchange step, at which point the effect of adding more GAP would saturate.

The maximum gain in rates for GAP-only and GEF-only assays is limited by the same basal GTP hydrolysis rate *r*_*h*,0_ and basal nucleotide exchange rate *r*_*e*,0_, leading to the following interdependence:

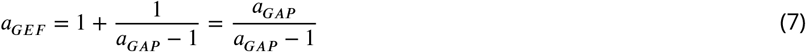

Analogously,

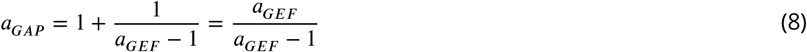

### Can the rate-limiting step model fully explain the experimental data? I.e. could rate-limiting steps be the *only* source of the positive *K*_3,*Cdc*24,*Rga*2_?

If the rate-limiting step model *fully* explains the synergy term observed in Cdc42 + GEF + GAP assays (Fig. 5), at least one of the two conditions, as already outlined in the section above, must be true:

1. the addition of a GEF speeds up the GDP release step (step 1) so much that the release step stops (or almost stops) being the rate-limiting step, or
2. the addition of a GAP speeds up the GTP hydrolysis step (step 2) so much that the hydrolysis step stops (or almost stops) being the rate-limiting step.

This ultimately means that the rate-limiting step model describes our data fully if at least one of the two the acceleration factors determined using our experimental data (*a*_*GEF*_ or *a*_*GAP*_) is consistent with the acceleration factor determined using the rate-limiting step model (Eq. 7 and 8).

### Acceleration factors from experimental data

The acceleration factors *a* describe the ratio of the basal GTPase cycle rate (i.e. the rate of Cdc42 alone, *r*_*c*,0_) over the GTPase cycle rate with one effector present (a GEF or a GAP) (*r*_*c*,1_) (Eq. 6). We determined the rates using an exponential fit

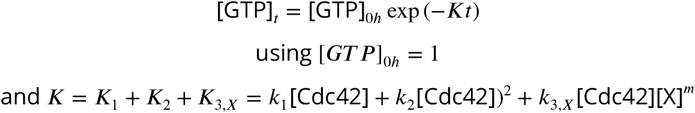

where *K* refers to the overall GTP hydrolysis rate and *X* is the effector protein (i.e. Cdc24, Rga2_(*I*)_, Rga2_(*II*)_, or the GAP domain). Without effectors present ([*X*] = 0) *K* = *r*_*c*,0_, and with an effector present *K* = *r*_*c*,1_.

*K* is concentration dependent and we here want to calculate it for maximal acceleration of the overall rate through GEF or GAP. Thus, we use 0.3 µM for Cdc24 (the highest Cdc24 concentration used in our assays) and 1.2 µM for all GAPs (given that the linear regime for both Rga2 versions extends up to 1.2 µM ^3^). Because the majority of assays were conducted using 1.3 µM Cdc42, we use 1.3 µM for Cdc42. Thus, the rates are:

- *K*_1_ + *K*_2_ = *k*_1_ [Cdc42] + *k*_2_ [Cdc42]^2^ = 0.165 h^−1^
- *K*_3,*Cdc*24_ = *k*_3,*Cdc*24_ [Cdc42] [Cdc24]^2^ = 1.391 h^−1^
- *K*_3,*Rga*2−*I*_ = *k*_3,*Rga*2−*I*_ [Cdc42] [Rga2_(*I*)_] = 0.186 h^−1^
- *K*_3,*Rga*2−*II*_ = *k*_3,*Rga*2−*II*_ [Cdc42] [Rga2_(*I I*)_] = 0.434 h^−1^
- *K*_3,*GAPdomain*_ = *r*_*c*,0_ + *k*_3,*GAPdomain*_ [Cdc42] [GAP domain] = 0.282 h^−1^

Using these rates, we can calculate the acceleration factors of GEF and GAP based on our experimental data (Eq. 6).

### Comparison of acceleration factors

We now compare the experimentally determined acceleration factors of GEF and GAP (using protein concentrations leading to maximal accelerations of the overall rate) with the acceleration factors determined using the rate-limiting step model (Eq. 7 and 8).

Thus, we check if the acceleration by the GEF limits the maximum acceleration by the GAP in the way predicted by the limiting step model:

1. We calculate the acceleration of the GEF based on our experimental data: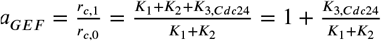.
2. We use Eq. 8 to calculate the maximum acceleration *a*_*GAP*_ that the GAP could achieve in the rate-limiting step model.
3. We compare *a*_*GAP*_ (rate-limiting step model) to *a*_*GAP*_ calculated from experimental data (e.g. for Rga2_(*I*)_: 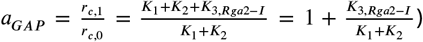. The rate-limiting step model describes the data fully if *a*_*GAP*_ (rate-limiting step model) ≥ *a*_*GAP*_ (experiment).

Analogously, we check if the acceleration by the GAP limits the maximum acceleration by the GEF in the way predicted by the rate-limiting step model:

1. We calculate the acceleration of the GAP based on our experimental data: e.g. for 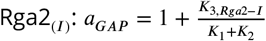.
2. We use Eq. 7 to calculate the maximum acceleration *a*_*GEF*_ that the GEF could achieve in the rate-limiting step model.
3. We compare *a*_*GEF*_ (rate-limiting step model) to *a*_*GEF*_ calculated from experimental data: 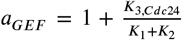. The rate-limiting step model describes the data fully if *a*_*GEF*_ (rate-limiting step model) ≥ *a*_*GEF*_ (experiment).

If one of the two aforementioned conditions on the rate-limiting effect of GAP/GEF addition is true, the rate-limiting step model describes our data fully and can be the sole source of the synergy term. The results are summarized in S5 Tab. 1: We determined a 9.43-fold increase the total rate from the Cdc42 + GEF assay data (experiment, Cdc42 + GEF). A *∼*10-fold acceleration factor of the GEF would maximize the GAP acceleration factor to 1.12 (rate-limiting step model, Cdc42 + GAP). For GAPs, we observe acceleration factors of 2.13 (Rga2_(*I*)_), 3.63 (Rga2_(*II*)_), and 2.71 (GAP domain) (experiment, GAP), which are all significantly larger than the by the rate-limiting step model predicted acceleration factor (i.e. *a*_*GAP*_ (rate-limiting step model) ≤ *a*_*GAP*_ (experiment)).

Similarly, the 2.13-fold (Rga2_(*I*)_), 3.63-fold (Rga2_(*II*)_), and 2.71-fold (GAP domain) increase the total rate from the Cdc42 + GAP assay data (experiment, Cdc42 + GAP) maximizes the GEF acceleration factor to 1.88 (Rga2_(*I*)_), 1.38 (Rga2_(*II*)_), and 1.58 (GAP domain) (rate-limiting step model, Cdc42 + GEF). For Cdc24, we observe an acceleration factor of 9.43 (experiment, Cdc42 + GEF). Here again, the experimentally observed rate acceleration by the GEF Cdc24 is much larger than the maximum acceleration predicted by the rate-limiting step model (i.e. *a*_*GEF*_ (rate-limiting step model) ≤ *a*_*GEF*_ (experiment)).

Thus, our experimental data does not fit the interdependence of the acceleration achieved by the GEF or GAP in the way described by the rate-limiting step model - the synergy term cannot be fully explained by the rate-limiting step model alone.

Furthermore, in Eq. 7 we assume that the GEF accelerates the cycle so much that the nucleotide exchange step is much faster than the hydrolysis step, at which point the effect of adding more GEF would saturate. We do not observe a GEF concentration regime where we see saturation. Thus in reality, the experimentally measured acceleration factor is likely an underestimation of the acceleration maximally possible, making the incompatibility between the rate-limiting step model and the acceleration factor data more pronounced.

Analogously, in Eq. 8 we assume that GAP accelerates the cycle so much that the hydrolysis step is much faster than the nucleotide exchange step, at which point the effect of adding more GAP would saturate. We do not observe a concentration regime for the GAP domain where we see saturation^4^. Thus, in reality the experimentally measured acceleration factor is likely an underestimation of the acceleration maximally possible.

**S5 Table 1.** Comparison of acceleration factors determined experimentally (Eq. 6) with those following the rate-limiting step model (Eq. 7, 8).

|  | Cdc42 + GEF<br>GEF = Cdc24 | Cdc42 + GAP |  |  |
| --- | --- | --- | --- | --- |
|  |  | GAP = Rga2<sub>(I)</sub> | GAP = Rga2<sub>(II)</sub> | GAP = GAP domain |
| **experiment:**<br>$a = \frac{r_{c,1}}{r_{c,0}} = 1 + \frac{K_{3,X}}{K_1+K_2}$ | $a_{GEF} = 1 + \frac{K_{3,Cdc24}}{K_1+K_2}$<br>= 9.43 | $a_{GAP} = 1 + \frac{K_{3,Rga2-I}}{K_1+K_2}$<br>= 2.13 | $a_{GAP} = 1 + \frac{K_{3,Rga2-II}}{K_1+K_2}$<br>= 3.63 | $a_{GAP} = 1 + \frac{K_{3,GAPdomain}}{K_1+K_2}$<br>= 2.71 |
| **rate-limiting step model:**<br>$a_{GEF} = \frac{a_{GAP}}{a_{GAP}-1}$<br>and<br>$a_{GAP} = \frac{a_{GEF}}{a_{GEF}-1}$ | GAP = Rga2<sub>(I)</sub>: $a_{GEF} = 1.88$<br>GAP = Rga2<sub>(II)</sub>: $a_{GEF} = 1.38$<br>GAP = GAP domain: $a_{GEF} = 1.58$ | $a_{GAP} = 1.12$ | | |

## Supplement S6

**S6 Figure 1.**
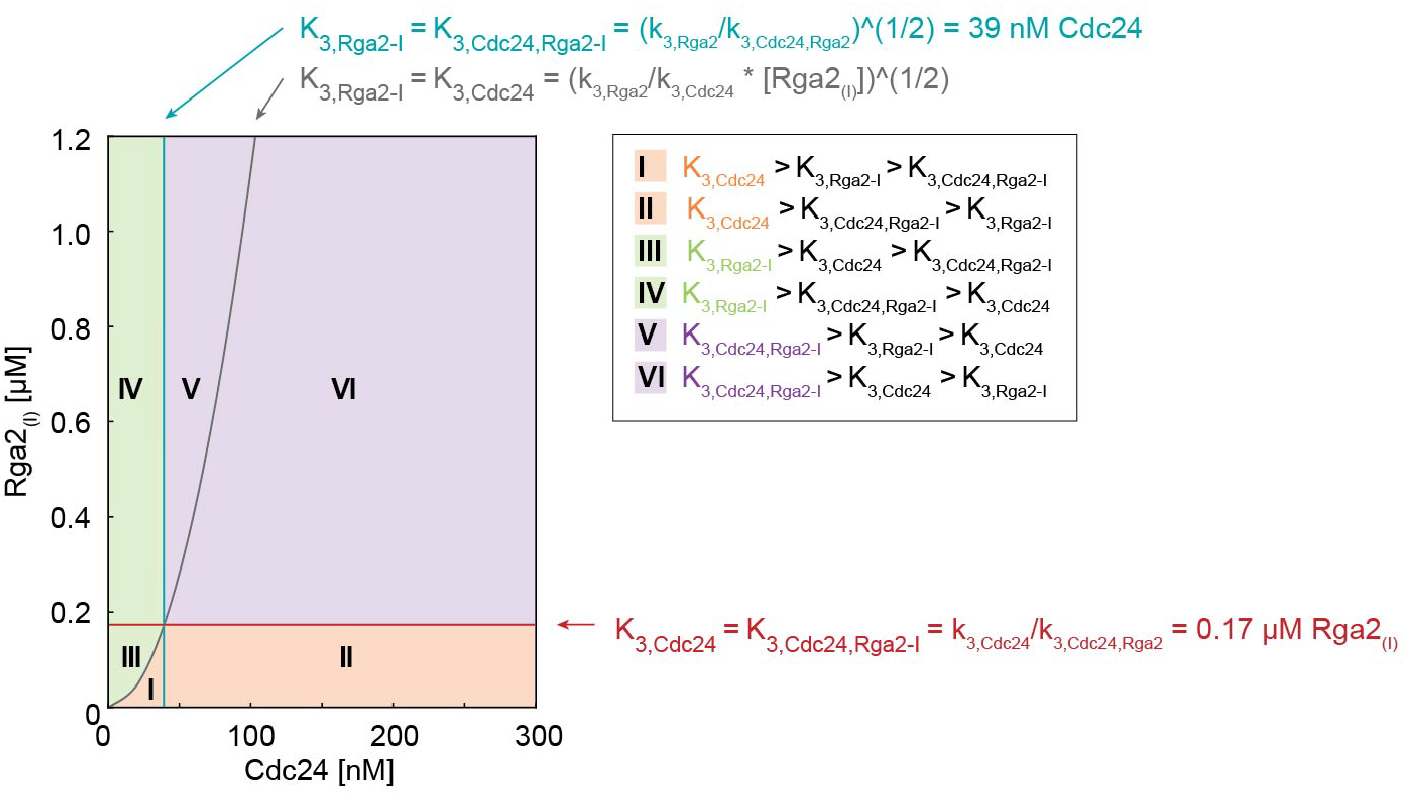
Regime diagram illustrating the dominant rate *K* across varying Cdc24 and Rga2_(*I*)_ concentrations. The diagram was generated using rates *k*_3_ for effectors Cdc24 and Rga2_(*I*)_ given in Tab. 3.

## Supplement S7

**S7 Figure 1.**
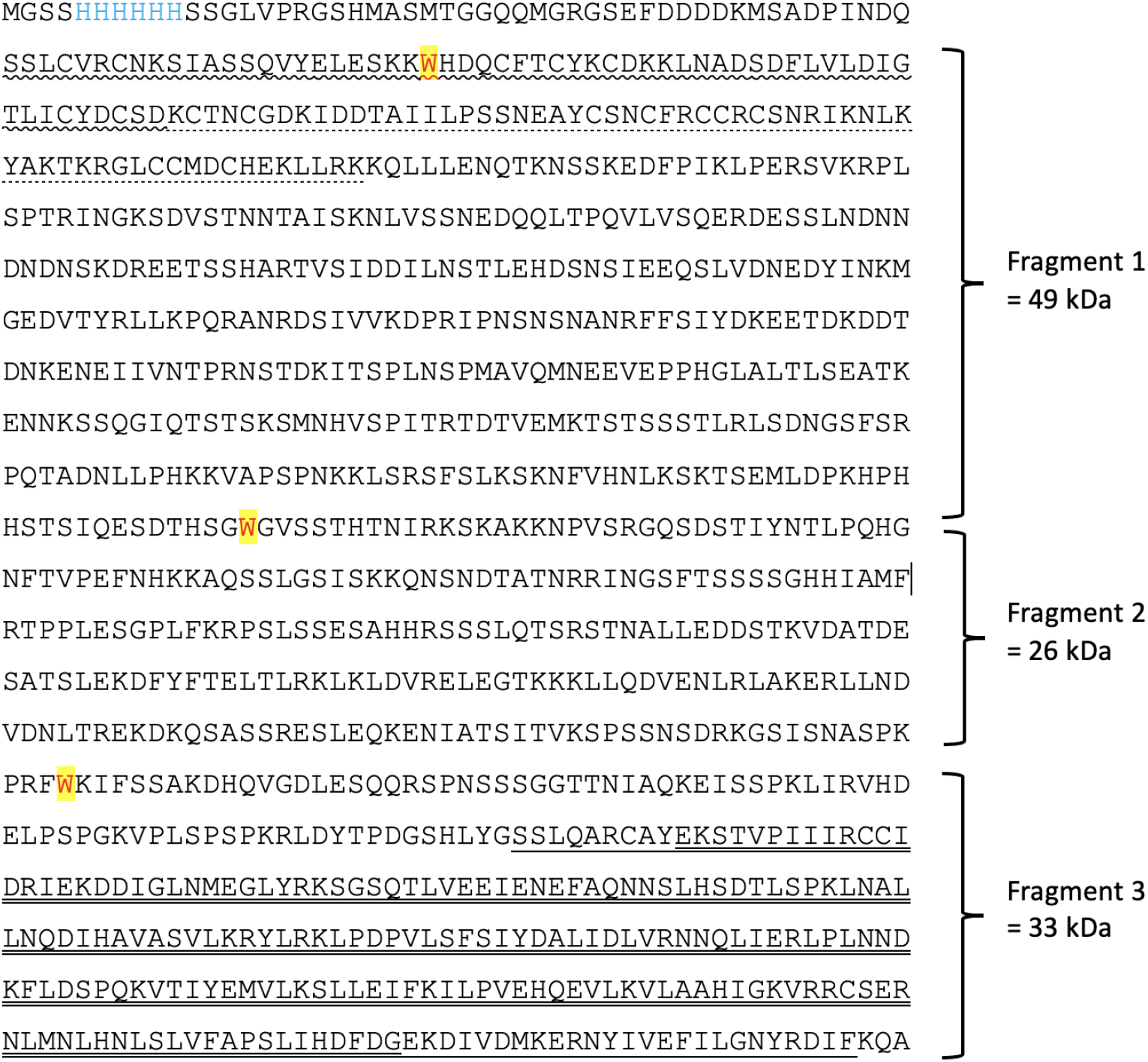
Amino acid sequence of the Rga2_(*I*)_ construct. Protein fragments that are not be visible on stain-free visualized SDS-PAGE (detection of tryptophan) or in anti-His Western blots are annotated on the right. The following sequence features are highlighted: 6His tag (blue), tryptophan (W) (red with yellow background), LIM zinc-binding domain (underlined, wriggled line), LIM zinc-binding domain (underlined, dotted line), RhoGAP domain (according to UNIPROT: underlined, single line; according to Smith *et al*. [Smith et al., 2002]: underlined, double line).

## Supplement S8

**S8 Figure 1.**
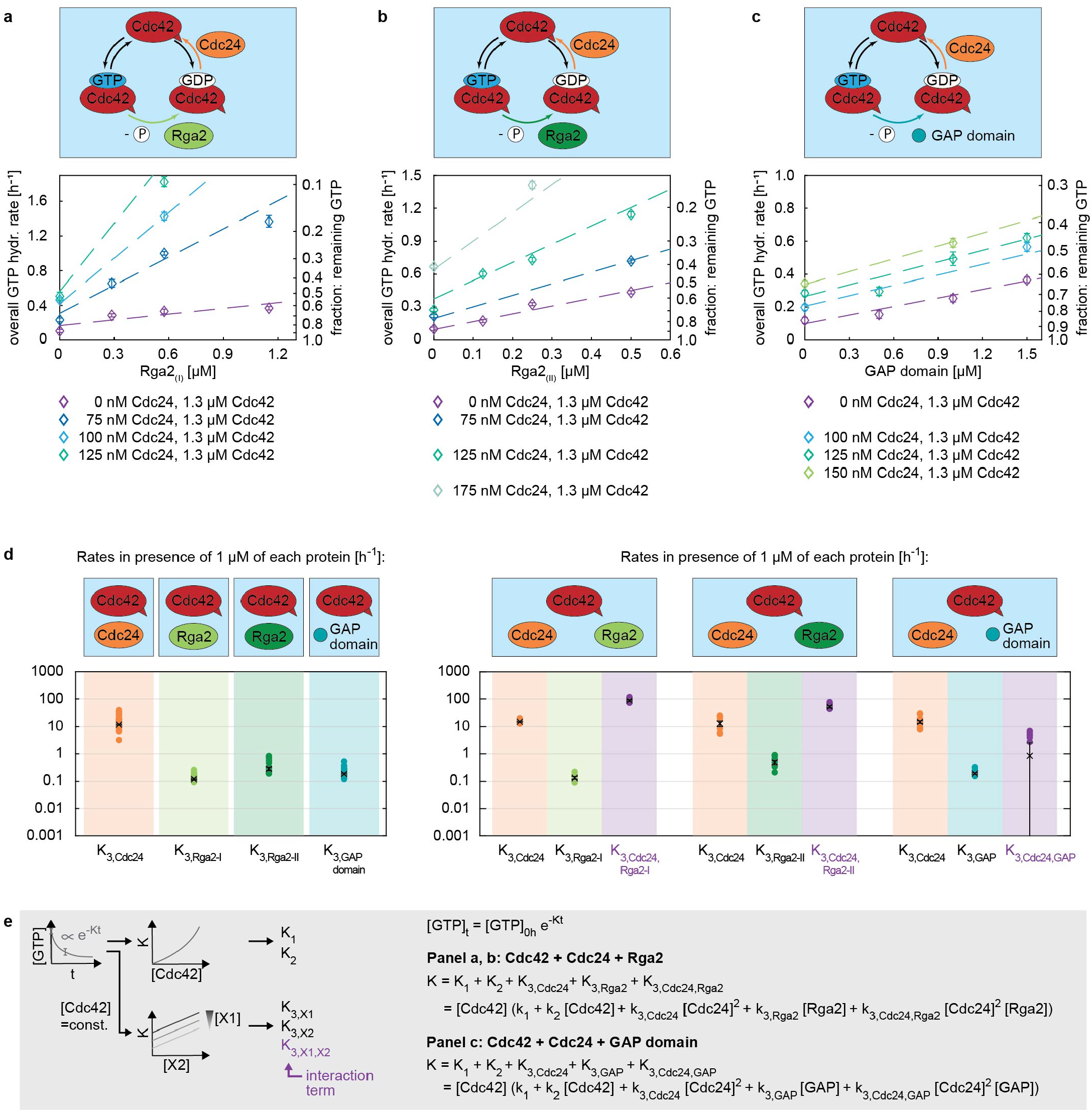
The C-terminal Flag-tag of Rga2_(*II*)_ might weaken Rga2-Cdc24 binding, reducing Cdc24-Rga2_(*II*)_ synergy. (a-c) GTPase assay data of (a) Cdc42-Cdc24-Rga2_(*I*)_, (b) Cdc42-Cdc24-Rga2_(*II*)_, and (c) Cdc42-Cdc24-GAP domain mixtures. (d) Summary of the rates *K* obtained in the three-protein assay (right) in comparison to those of the two-protein assay (left): In three-protein assays the rate contribution of the individual proteins is comparable to those obtained in the two-protein assay. Additionally, an interaction rate is obtained (shown in purple). Both Cdc42-Cdc24-Rga2_(*I*)_ and Cdc42-Cdc24-Rga2_(*I*)_ show a positive interaction rate, with *K*_3,*Cdc*24,*Rga*2−*I*_ > *K*_3,*Cdc*24,*Rga*2−*II*_ (Tab. 3). The C-terminal Flag-tag of Rga2_(*II*)_ might weaken Cdc24-Rga2 binding, reducing Cdc24-Rga2_(*II*)_ synergy. The values shown refer to the rate values in presence of 1 µM of each protein, e.g. ‘*K*_3,*Cdc*24_’ refers to ‘*k*_3,*Cdc*24_ [Cdc24]^2^ [Cdc42]’ with [Cdc42]=[Cdc24]=1 µM. Crosses with error bars represent the weighted mean and the standard error of the mean (S1), and filled circles show individual measurements. (e) Illustration of the data processing and fitting model.

## Supplement S9

**S9 Figure 1.**
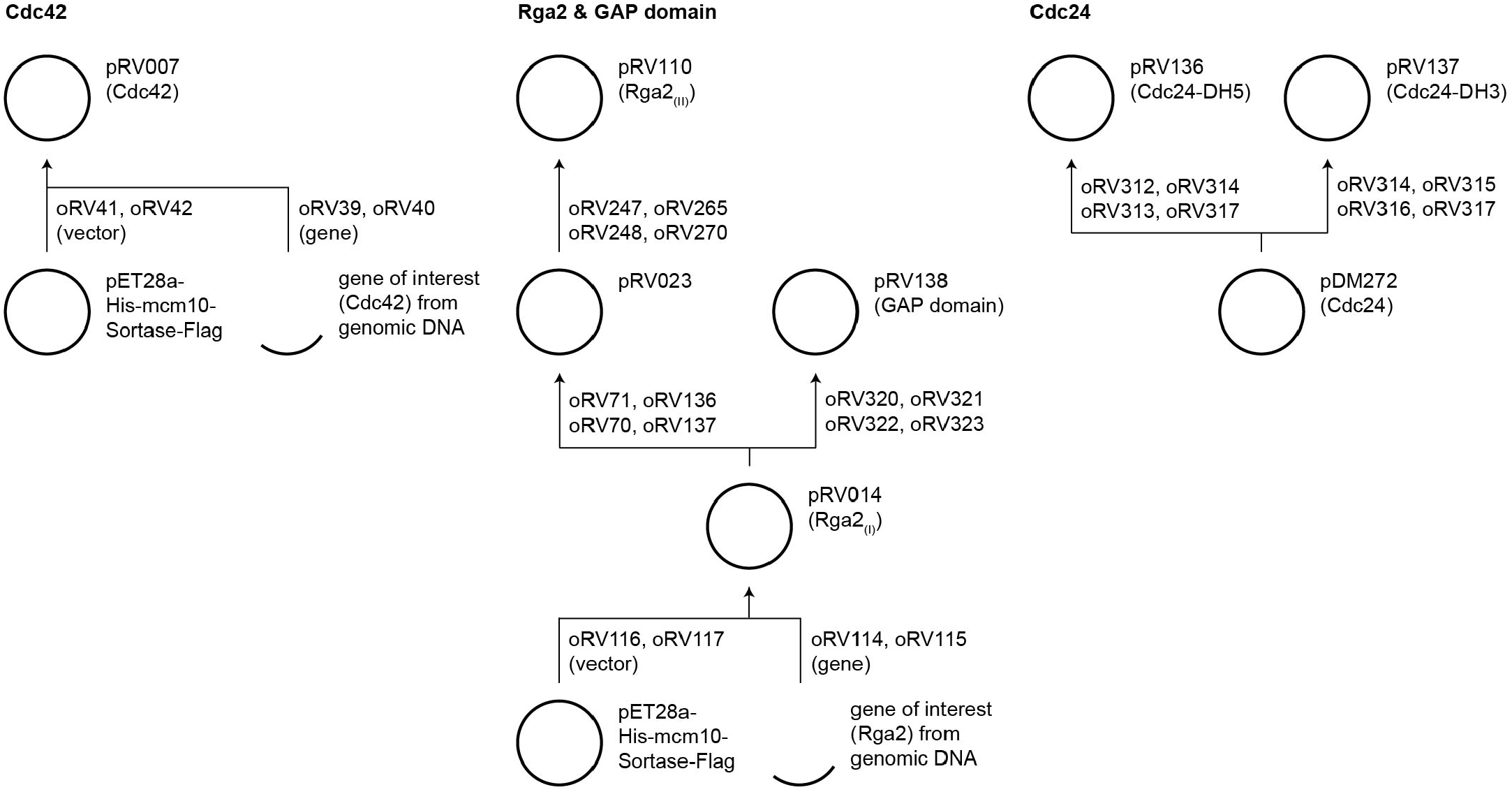
Schematic of the plasmid construction. pET28a-His-mcm10-Sortase-Flag was received from N. Dekker (TU Delft) and is based on pBP6 [Douglas and Diffley, 2016]. pDM272 was received from D. McCusker (University of Bordeaux) [Rapali et al., 2017].

**S9 Table 1.**
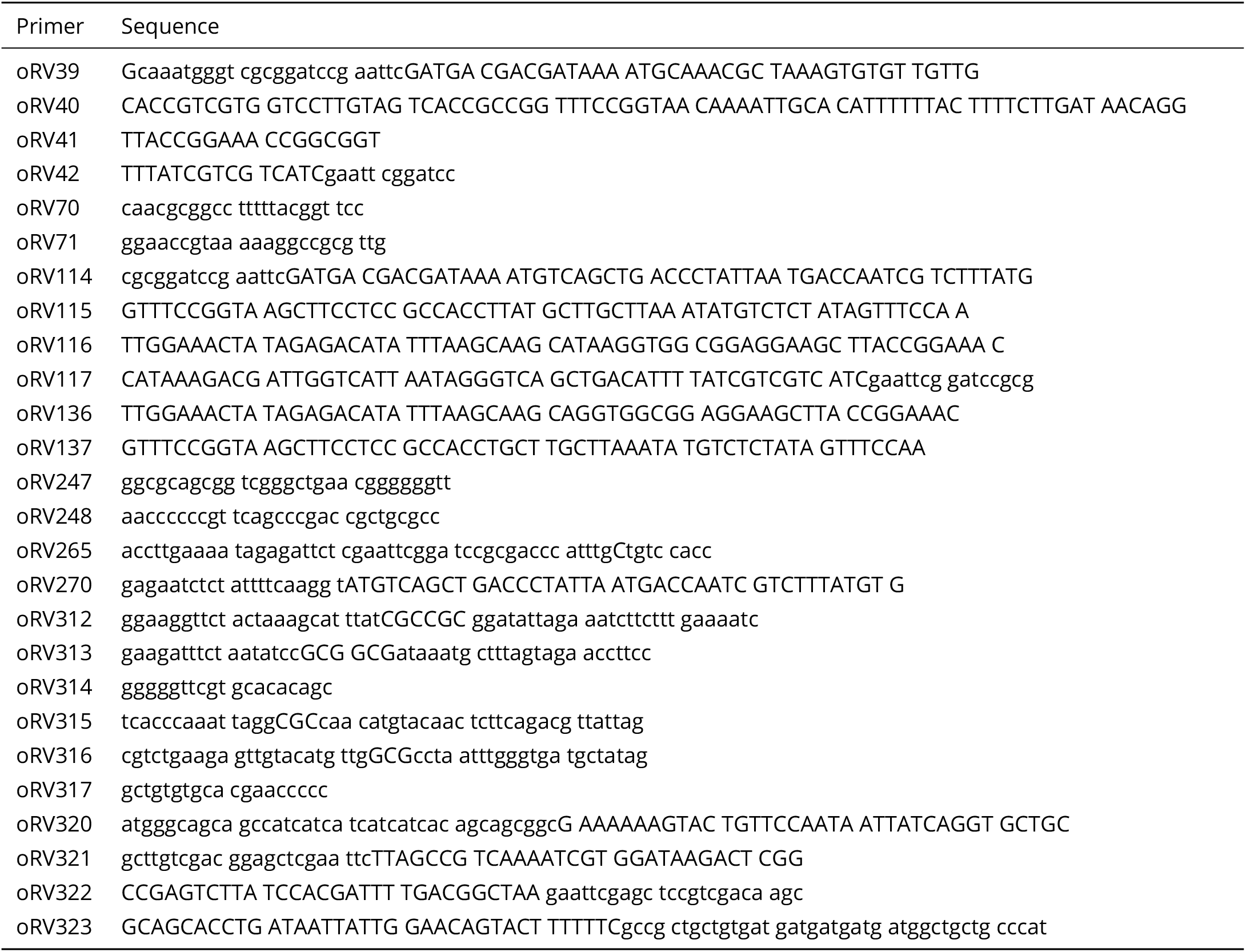
Primer overview.

## Supplement S10

### Cdc42, pRV007

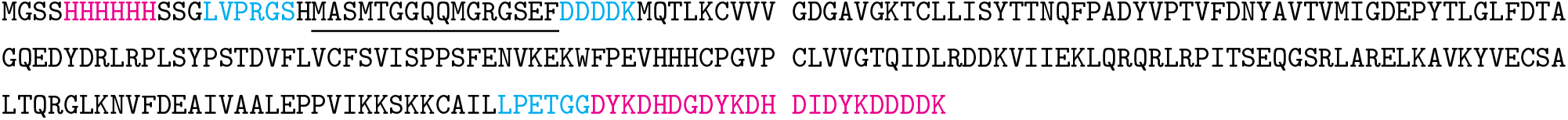

### Rga2_(*I*)_, pRV014

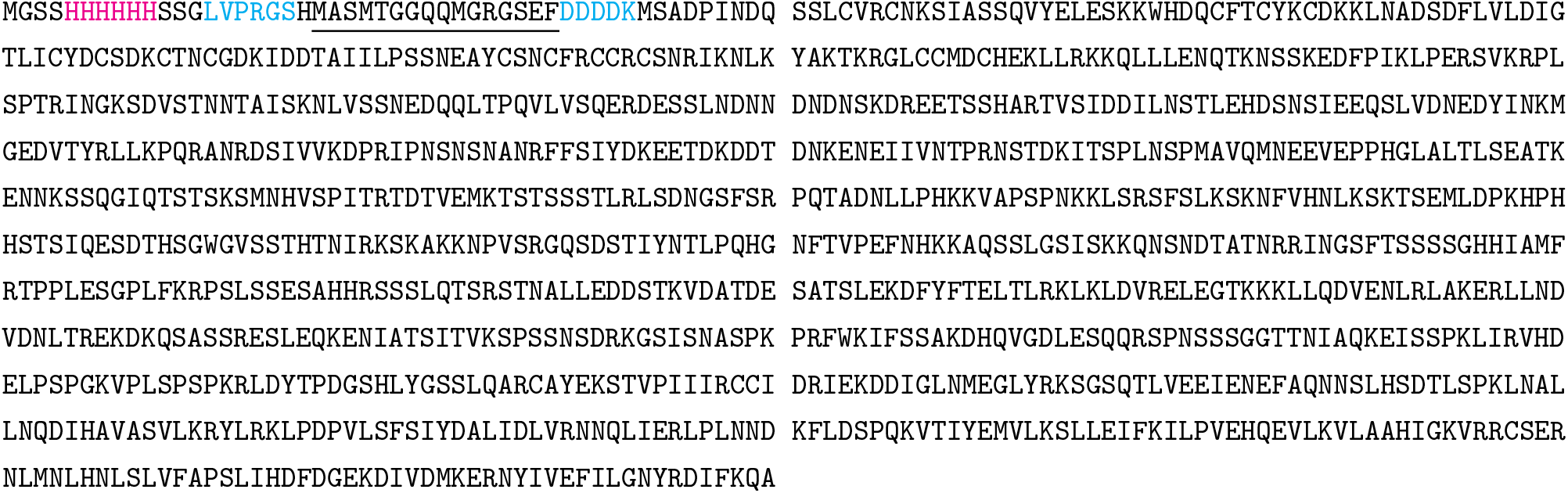

### Rga2_(*II*)_, pRV110

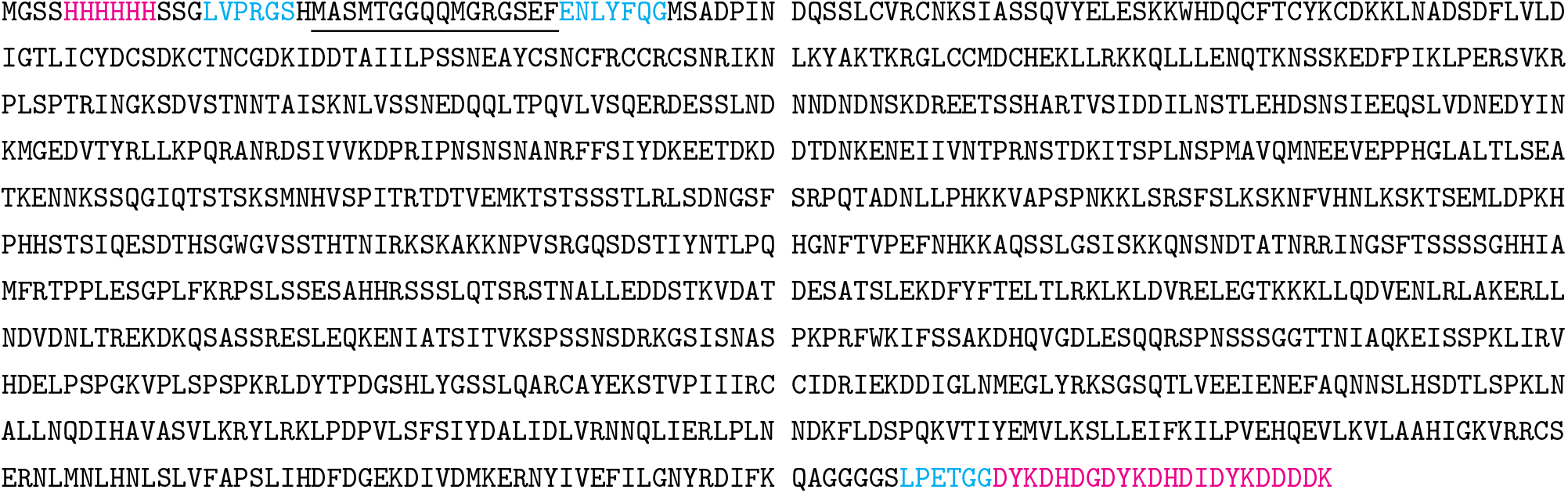

### GAP domain (amino acids 797-981 from Rga2 [Smith et al., 2002]), pRV138

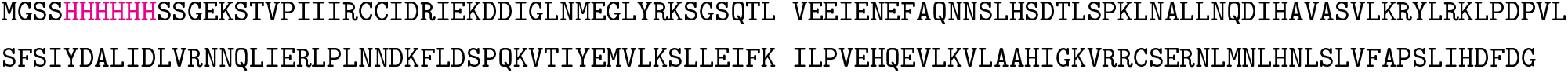

### Cdc24, pDM272

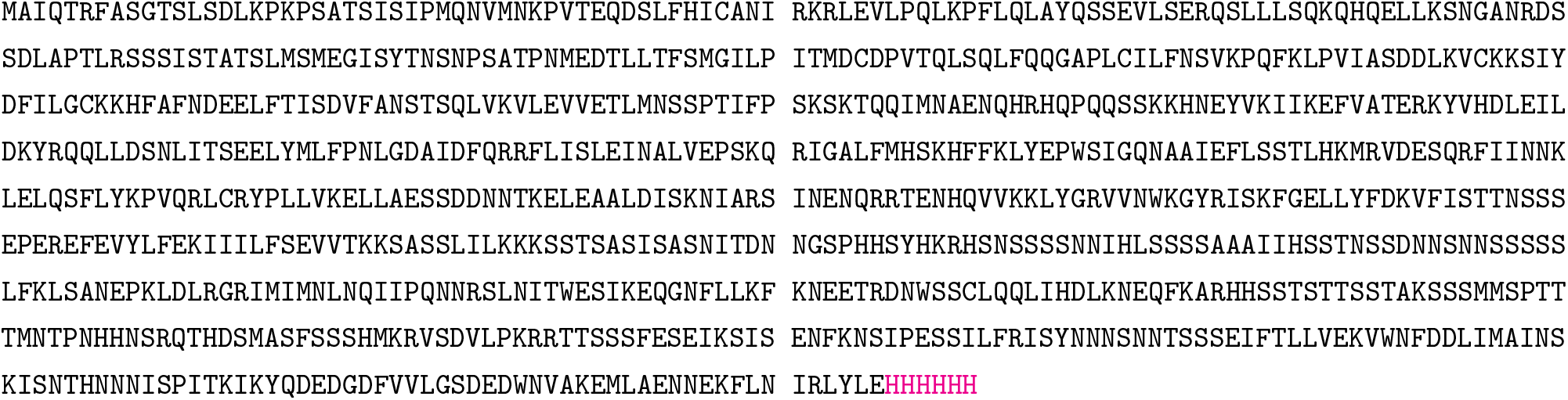

### Cdc24-DH5 (Cdc24 mutation F322A [Mionnet et al., 2008]), pRV136

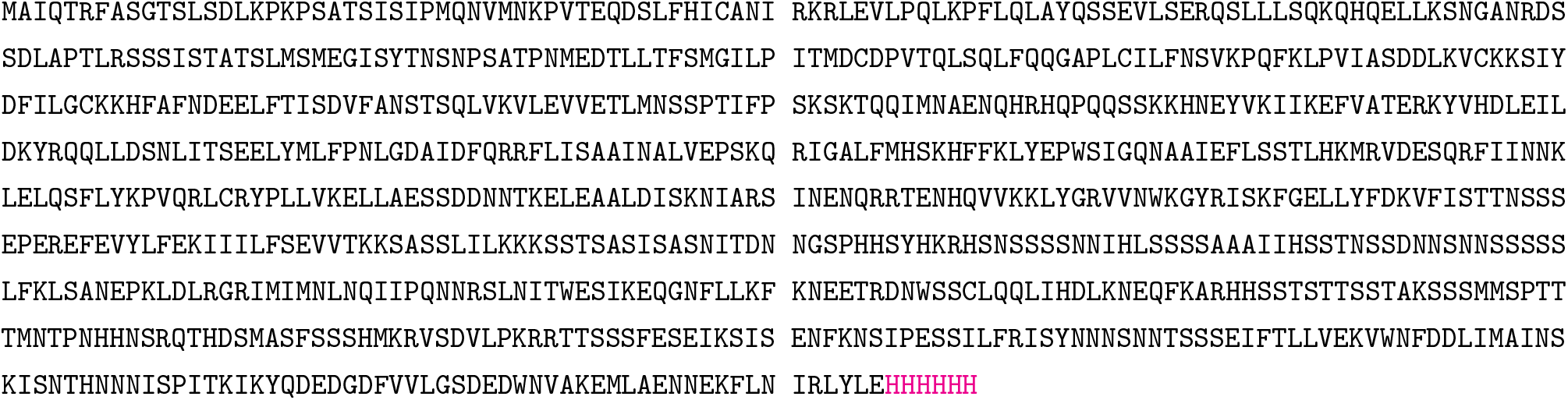

### Cdc24-DH3 (Cdc24 mutations L339A and E340A [Mionnet et al., 2008]), pRV137

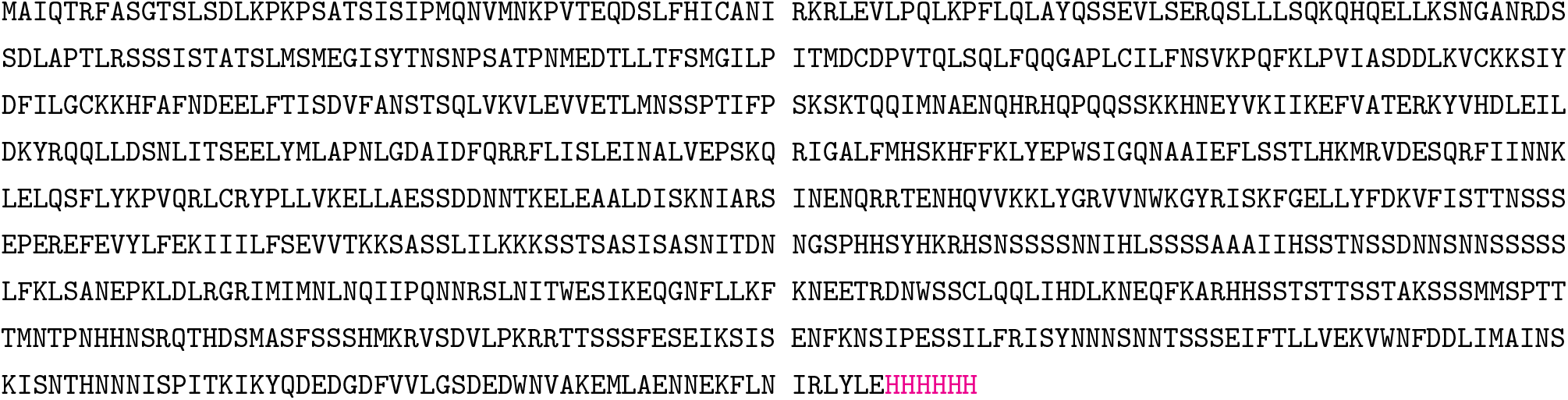

The protein constructs contain some of the following features:

1. 6His-tag: HHHHHH
2. Flag-tag: DYKDHDGDYKDHDIDYKDDDDK
3. Thrombin cut site: LVPRGS
4. Enterokinase cut site: DDDDK
5. TEV cut site: ENLYFQG
6. Sortase cut/ligation site: LPETGG
7. T7 tag (to aid protein expression): <u>MASMTGGQQMGRGSEF</u>

More information on the protein constructs is given in [Tschirpke et al., 2023b].

## Supplement S11

**S11 Figure 1.**
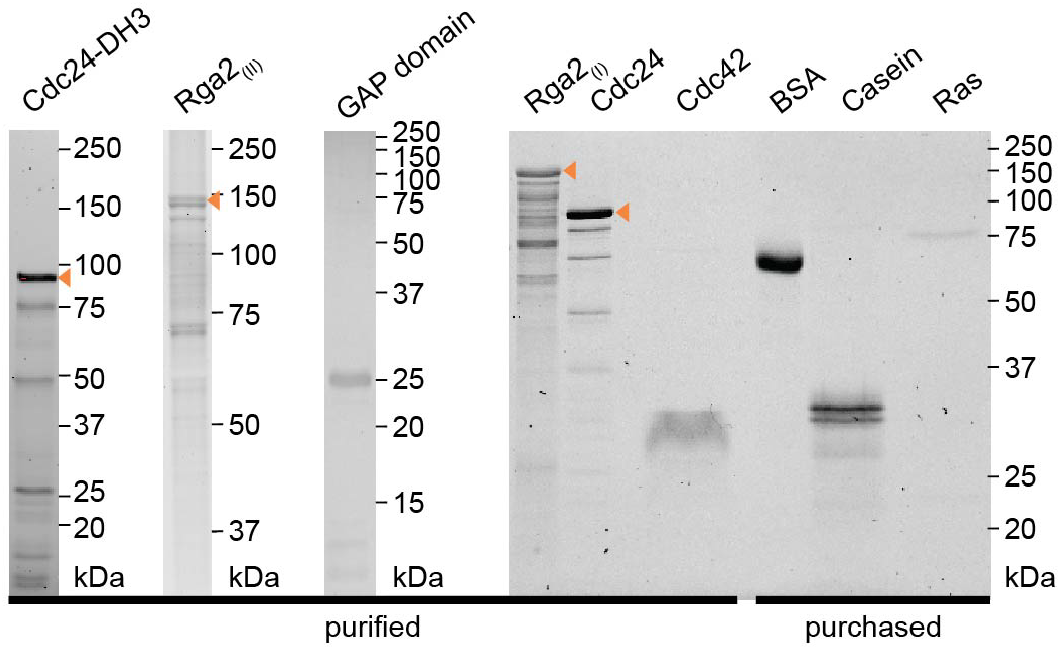
SDS-PAGE of used proteins. An orange arrow indicates the band of the correct size. The GAP domain is visualized using SimplyBlue SafeStain while all other gels show data obtained from stain-free imaging

## Footnotes

1 We attempted to reproduce Cdc24 oligomerization using SEC-MALS experiments, but did not observe Cdc24 oligomers (S3). Given that Cdc24 oligomerizes via weak interactions [Mionnet et al., 2008] and that SEC-MALS experiments are performed under a constant flow that could pull weakly bound Cdc24 oligomers apart, our results do not contradict previous findings.

2 The fit quality for the Cdc42-Rga2-BSA data was significantly lower than for all other data sets (*R*^2^ ≈ 0.5). Hence, we remain cautious drawing conclusions from this data.

3 We observed a linear regime for the GAP domain also above 1.2 µM. To keep the comparison for all GAPs consistent, we still use 1.2 µM for the GAP domain.

4 We observe saturation for Rga2, but believe that this saturation is due to auto-inhibition of Rga2. If this saturation where because the hydrolysis step cannot go faster/ is reaching a rate-limiting step, we would also observe saturation for the GAP domain (which we do not).

## References

Breitkreutz, A., et al. (2010). A global protein kinase and phosphatase interaction network in yeast. Science, 328(5981), 1043–1046, 10.1126/science.1176495.

Chebotareva, N. A., Kurganov, B. I., & Livanova, N. B. (2004). Biochemical Effects of Molecular Crowding. Biochemistry (Moscow), 69(11), 1239–1251.

Chiou, J.-g., Balasubramanian, M. K., & Lew, D. J. (2017). Cell Polarity in Yeast. Annual Review of Cell and Developmental Biology, 33, 77–101, 10.1146/annurev.cellbio.15.1.365 https://doi.org/10.1146/annurev-cellbio-100616-060856.

Chollet, J., Dunkler, A., Bauerle, A., Vivero-Pol, L., Mulaw, M. A., Gronemeyer, T., & Johnsson, N. (2020). Cdc24 interacts with septins to create a positive feedback loop during bud site assembly in yeast. Journal of Cell Science, 133(11), 10.1242/jcs.240283.

Daalman, W., Sweep, E., & Laan, L. (2020). The Path towards Predicting Evolution as Illustrated in Yeast Cell Polarity. Cells, 9(12), 2534, 10.3390/cells9122534 https://www.mdpi.com/2073-4409/9/12/2534.

Das, A., Slaughter, B. D., Unruh, J. R., Bradford, W. D., Alexander, R., Rubinstein, B., & Li, R. (2012). Flippase-mediated phospholipid asymmetry promotes fast Cdc42 recycling in dynamic maintenance of cell polarity. Nature Cell Biology, 14(3), 304–310, 10.1038/ncb2444 http://www.nature.com/doifinder/10.1038/ncb2444.

Diepeveen, E. T., Gehrmann, T., Pourquié, V., Abeel, T., & Laan, L. (2018). Patterns of conservation and diversification in the fungal polarization network. Genome Biology and Evolution, 10(August), evy121–evy121, 10.1093/gbe/evy121 http://dx.doi.org/10.1093/gbe/evy121.

Douglas, M. E. & Diffley, J. F. (2016). Recruitment of Mcm10 to sites of replication initiation requires direct binding to the minichromosome maintenance (MCM) complex. Journal of Biological Chemistry, 291(11), 5879–5888, 10.1074/jbc.M115.707802.

Etienne-Manneville, S. (2004). Cdc42–the centre of polarity. J Cell Sci, 117(Pt 8), 1291–1300, 10.1242/jcs.01115 http://www.ncbi.nlm.nih.gov/pubmed/15020669.

Gao, J. T., et al. (2011). Modular coherence of protein dynamics in yeast cell polarity system. Proceedings of the National Academy of Sciences, 108(18), 7647–7652, 10.1073/pnas.1017567108 http://www.pnas.org/cgi/doi/10.1073/pnas.1017567108.

Gibson, D. G., Young, L., Chuang, R. Y., Venter, J. C., Hutchison, C. A., & Smith, H. O. (2009). Enzymatic assembly of DNA molecules up to several hundred kilobases. Nature Methods, 6(5), 343–345, 10.1038/nmeth.1318.

Golding, A. E., Visco, I., Bieling, P., & Bement, W. M. (2019). Extraction of active RhoGTPases by RhoGDI regulates spatiotemporal patterning of RhoGTPases. eLife, 8, e50471, 10.7554/eLife.50471 https://elifesciences.org/articles/50471.

Heij, C., de Boer, P., Franses, P. H., Kloek, T., & van Dijk, H. K. (2004). Econometric Methods with Applications in Business and Economics. Oxford University Press.

Johnson, J. L., Erickson, J. W., & Cerione, R. A. (2009). New Insights into How the Rho Guanine Nucleotide Dissociation Inhibitor Regulates the Interaction of Cdc42 with Membranes. Journal of Biological Chemistry, 284(35), 23860– 23871, 10.1074/jbc.M109.031815 http://www.ncbi.nlm.nih.gov/pubmed/19581296 whttps://linkinghub.elsevier.com/retrieve/pii/S0021925818608130.

Johnson, J. L., Erickson, J. W., & Cerione, R. A. (2012). C-terminal Di-arginine motif of Cdc42 protein is es-sential for binding to phosphatidylinositol 4,5-bisphosphate-containing membranes and inducing cellular transformation. Journal of Biological Chemistry, 287(8), 5764–5774, 10.1074/jbc.M111.336487.

Kang, P. J., Béven, L., Hariharan, S., & Park, H. O. (2010). The Rsr1/Bud1 GTPase interacts with itself and the Cdc42 GTPase during bud-site selection and polarity establishment in budding yeast. Molecular Biology of the Cell, 21(17), 3007–3016, 10.1091/mbc.E10-03-0232.

Laemmli, U. K. (1970). Cleavage of structural proteins during the assembly of the head of bacteriophage T4. Nature, 227, 680–685 https://doi-org.tudelft.idm.oclc.org/10.1038/227680a0.

Lang, C. F. & Munro, E. M. (2022). Oligomerization of peripheral membrane proteins provides tunable control of cell surface polarity. Biophysical Journal, 121(23), 4543–4559, 10.1016/j.bpj.2022.10.035 https://doi.org/10.1016/j.bpj.2022.10.035.

Lee, C.-H., Occhipinti, P., & Gladfelter, A. S. (2015). PolyQ-dependent RNA-protein assemblies control symmetry breaking. Journal of Cell Biology, 208(5), 533–544, 10.1083/jcb.201407105.

Martin, S. G. (2015). Spontaneous cell polarization: Feedback control of Cdc42 GTPase breaks cellular symmetry. BioEssays, 37(11), 1193–1201, 10.1002/bies.201500077.

McCusker, D., Denison, C., Anderson, S., Egelhofer, T. A., Yates, J. R., Gygi, S. P., & Kellogg, D. R. (2007). Cdk1 coordinates cell-surface growth with the cell cycle. Nature Cell Biology, 9(5), 506–515, 10.1038/ncb1568.

Mionnet, C., Bogliolo, S., & Arkowitz, R. A. (2008). Oligomerization regulates the localization of Cdc24, the Cdc42 activator in Saccharomyces cerevisiae. Journal of Biological Chemistry, 283(25), 17515–17530, 10.1074/jbc.M800305200.

Nelson, W. J. (2003). Generating polarity in single cells for mitotic division. Nature, 433(April), 766–774 www.nature.com/nature.

Park, H.-O. & Bi, E. (2007). Central Roles of Small GTPases in the Development of Cell Polarity in Yeast and Beyond. Microbiology and Molecular Biology Reviews, 71(1), 48–96, 10.1128/MMBR.00028-06 http://mmbr. asm.org/cgi/doi/10.1128/MMBR.00028-06.

Rapali, P., et al. (2017). Scaffold-mediated gating of Cdc42 signalling flux. eLife, 6, 1–18, https://doi.org/10.7554/eLife.25257 whttps://elifesciences.org/articles/25257.

Ross, E. M. (2008). Coordinating Speed and Amplitude in G-Protein Signaling. Current Biology, 18(17), 10.1016/j.cub.2008.07.035.

Sartorel, E., Unlu, C., Jose, M., Aurélie, M. L., Meca, J., Sibarita, J. B., & McCusker, D. (2018). Phosphatidylserine and GTPase activation control Cdc42 nanoclustering to counter dissipative diffusion. Molecular Biology of the Cell, 29(11), 1299–1310, 10.1091/mbc.E18-01-0051.

Schlecht, U., Miranda, M., Suresh, S., Davis, R. W., & St. Onge, R. P. (2012). Multiplex assay for conditiondependent changes in protein-protein interactions. Proceedings of the National Academy of Sciences of the United States of America, 109(23), 9213–9218, 10.1073/pnas.1204952109.

Shimada, Y., Gulli, M.-P., & Peter, M. (2000). Nuclear sequestration of the exchange factor Cdc24p by Far1 regulates cell polarity during mating. Nature Cell Biology, 2, 117–124 https://doi-org.tudelft.idm.oclc.org/10.1038/35000073.

Shimada, Y., Wiget, P., Gulli, M. P., Bi, E., & Peter, M. (2004). The nucleotide exchange factor Cdc24p may be regulated by auto-inhibition. EMBO Journal, 23(5), 1051–1062, 10.1038/sj.emboj.7600124.

Smith, G. R., Givan, S. A., Cullen, P., & Sprague, G. F. (2002). GTPase-Activating Proteins for Cdc42. Eukaryotic Cell, 1(3), 469–480, 10.1128/EC.1.3.469-480.2002 https://journals.asm.org/doi/10.1128/EC.1.3.469-480.2002.

Smith, S. E., et al. (2013). Independence of symmetry breaking on Bem1-mediated autocatalytic activation of Cdc42. Journal of Cell Biology, 202(7), 1091–1106, 10.1083/jcb.201304180 http://www.ncbi.nlm.nih.gov/pubmed/24062340.

Tarassov, K., et al. (2008). An in vivo map of the yeast protein interactome. Science, 320(5882), 1465– 1470, 10.1126/science.1153878.

Thompson, B. J. (2013). Cell polarity: Models and mechanisms from yeast, worms and flies. Development (Cambridge), 140(1), 13–21, 10.1242/dev.083634.

Tschirpke, S., Daalman, W. K., & Laan, L. (2024). Quantification of GTPase Cycling Rates of GTPases and GTPase:Effector Mixtures Using GTPase Glo Assays. Current Protocols, 4(4), 1–29, 10.1002/cpz1.1000.

Tschirpke, S., Daalman, W. K. G., & Laan, L. (2023a). Data accompanying the publication: Quantification of GTPase cycling rates of GTPases and GTPase: effector mixtures using GTPase Glo assays. 4TU.ResearchData (CC BY-SA 4.0), 10.4121/ac196f25-1c20-4c0c-a0b9-f01cd3fadc45.

Tschirpke, S., Daalman, W. K. G., & Laan, L. (2025). Data underlying the publication: Oligomerizationdependent and synergistic regulation of Cdc42 GTPase cycling by a GEF and a GAP. 4TU.ResearchData (CC BY-SA 4.0), 10.4121/56535d83-8a7a-4367-9ae5-f0d56b51161f.

Tschirpke, S., van Opstal, F., van der Valk, R., Daalman, W. K.-G., & Laan, L. (2023b). A guide to the in vitro reconstitution of Cdc42 activity and its regulation. BioRxiv, 10.1101/2023.04.24.538075.

Vendel, K. J. A., Tschirpke, S., Shamsi, F., Dogterom, M., & Laan, L. (2019). Minimal in vitro systems shed light on cell polarity. Journal of Cell Science, 132(4), jcs217554, 10.1242/jcs.21755410.1242/jcs.217554.

Wedlich-Soldner, R., Wai, S. C., Schmidt, T., & Li, R. (2004). Robust cell polarity is a dynamic state established by coupling transport and GTPase signaling. Journal of Cell Biology, 166(6), 889–900, 10.1083/jcb.200405061 https://rupress.org/jcb/article/166/6/889/51279/Robust-cell-polarity-is-a-dynamic-state.

Zhang, B., Gao, Y., Moon, S. Y., Zhang, Y., & Zheng, Y. (2001). Oligomerization of Rac1 GTPase Mediated by the Carboxyl-terminal Polybasic Domain. Journal of Biological Chemistry, 276(12), 8958–8967, 10.1074/jbc.M008720200 http://dx.doi.org/10.1074/jbc.M008720200.

Zhang, B., Wang, Z. X., & Zheng, Y. (1997). Characterization of the interactions between the small GTPase Cdc42 and its GTPase-activating proteins and putative effectors: Comparison of kinetic properties of Cdc42 binding to the Cdc42-interactive domains. Journal of Biological Chemistry, 272(35), 21999–22007, 10.1074/jbc.272.35.21999.

Zhang, B., Zhang, Y., Collins, C. C., Johnson, D. I., & Zheng, Y. (1999). A built-in arginine finger triggers the self-stimulatory GTPase-activating activity of Rho family GTPases. Journal of Biological Chemistry, 274(5), 2609–2612, 10.1074/jbc.274.5.2609.

Zhang, B. & Zheng, Y. (1998). Negative regulation of Rho family GTPases Cdc42 and Rac2 by homodimer formation. Journal of Biological Chemistry, 273(40), 25728–25733, 10.1074/jbc.273.40.25728.

Zheng, Y., Bender, A., & Cerione, R. A. (1995). Interactions among proteins involved in bud-site selection and bud-site assembly in Saccharomyces cerevisiae. Journal of Biological Chemistry, 270(2), 626–630 http://www.ncbi.nlm. nih.gov/pubmed/7822288.

Zheng, Y., Cerione, R., & Bender, A. (1994). Control of the yeast bud-site assembly GTPase Cdc42. Catalysis of guanine nucleotide exchange by Cdc24 and stimulation of GTPase activity by Bem3. J Biol Chem, 269(4), 2369–2372 http://www.ncbi.nlm.nih.gov/pubmed/8300560.

Zheng, Y., Harts, M. J., Shinjosq, K., Evansn, T., Benderll, A., & Ceriones, R. a. (1993). Biochemical Comparisons of the Saccharomyces cerevisiae Bem2 and Bem3 Proteins. Journal of Biological Chemistry, 268(33), 24629–24634 10.1016/S0021-9258(19)74512-8.

